# Using insertable cardiac monitors to test determinants of heart rate and activity in captive baboons

**DOI:** 10.64898/2026.03.13.710869

**Authors:** Catherine R. Andreadis, Ipek G. Kulahci, James Ndung’u, Daisy Kigen, Priscilla Kimiti, Kevin Mugambi Kibe, Noelle R. Laske, Janet Mwadime, Nashon Wanjala, Herman Pontzer, Timothy G. Laske, Mercy Y. Akinyi, Elizabeth A. Archie

## Abstract

**Background:** Insertable cardiac monitors (ICMs) provide fine-grained, continuous data on cardiac activity. These data have great potential to reveal individual physiology, energetics, and stress responses, with implications for animal health, cognition, welfare, and conservation. However, these devices must be tested for safety, accuracy, and biological validity before being deployed in new species. Here we do so for the Reveal LINQ^TM^ ICM (Medtronic, Minneapolis, MN USA) over an 8-month period in 10 adult female baboons (*Papio anubis* and *P. cynocephalus*) at the Kenya Institute of Primate Research in Kenya. We also report data on heart rate, physical activity, and body temperature in unrestrained, conscious, captive baboons during their normal activities. Finally, we test how heart rate and activity levels are predicted by baboon species, body mass, time of day, ambient temperature, social dominance rank, and ovarian cycle phase.

**Results:** The baboons had no adverse reactions to the Reveal LINQ^TM^ ICM. Their mean daytime heart rates (HRs) ranged from 89.6 to 128.0 beats per minute (bpm), and their resting HRs ranged from 74.4 to 98.2 bpm. The fastest observed R-wave interval validated by electrocardiogram (ECG) was 230 milliseconds (ms) (260 bpm), and the slowest was 1270 ms (47.4 bpm). In terms of predictors of HR and activity, HR was highly individualized, while activity level was not: baboon identity explained 41% of the variation in HR, but identity only 2% of variation in activity levels. HR was positively correlated with physical activity and HR was highest during daylight hours when the baboons were more active. Dominant baboons had higher HRs controlling for activity and were more active than low ranking individuals. In terms of ovarian cycle phase, HR was higher when individuals were in the periovulatory and luteal phases of the ovarian cycle compared to the follicular phase.

**Conclusions:** Our findings support the future use of ICMs to investigate physiological responses in baboons. These devices’ safety and validity represent the plausibility of understanding inter-individual and inter-species variation in heart rate and activity in response to variation in the external environment and in individual internal state.

## BACKGROUND

To survive and thrive, animals must respond to changes in their environment in ways that are both physiologically appropriate and return them to a state of homeostasis (1–4). Understanding these changes requires fine-grained, temporally resolved data on individual physiology as animals experience fluctuating weather, resources, threats, and other aspects of the environment. Such data are challenging to collect, but recent advances in physiologger technology are helping close the gap (5,6). For instance, physiologgers have recently shed light on individual feeding behavior and physiological responses to biological stressors such as parasitism (7,8). Physiologgers are also providing new insights into difficult to study physiological states, such as sleep and hibernation (9–13). In terms of conservation, physiologgers are revealing valuable information on the physiology of rare species, with implications for animal welfare and management (14).

One area where physiologgers are making major advances is in measuring real-time changes in animals’ heart rate (5,15,16). Heart rate physiologgers come in many forms, including collars, vests, belts, and internal devices that are either swallowed or surgically implanted (17–21). One of the most common and reliable such devices are subcutaneous implants known as insertable cardiac monitors (ICMs), which measure cardiac activity via subcutaneous electrical signals. Their use in wild animals is especially promising, allowing researchers to measure individual heart rate on minute-to-minute scales, infer metabolic rates, and to estimate the energetic costs of specific behaviors and life history traits (22–24). When viewed collectively, studies that leverage data from ICMs provide unprecedented insight into comparative patterns of cardiac physiology in many vertebrate species (10,14,23–31). ICMs are also useful for assessing aspects of the autonomic stress response via the sympathetic–adrenal–medullary system’s effects on heart rate (15). For example, heart rate data from ICMs has provided insight into how individuals perceive anthropogenic stressors such as roads, drones, or human encounters (28–30,32–35), as well as stress during capture, restraint, and transport of endangered species (31).

Despite their potential value, ICMs must be tested for accuracy, safety, and biological validity before being used in new species. Here, we tested these attributes for the Reveal LINQ^TM^ ICM (Medtronic, Minneapolis, MN USA) in captive baboons housed at the Kenya Institute of Primate Research (KIPRE), in Nairobi, Kenya. This work was a first step to ultimately deploying these ICMs in wild baboons. The Reveal LINQ^TM^ was developed for human clinical use, and differences human and baboon cardiac and musculoskeletal anatomy could cause variation in electrical noise (36). This could result in the misidentification of electrical noise as spurious heart beats when used in nonhuman species, such as baboons. However, this ICM has already been used successfully in several other mammalian species such as scimitar-horned oryx (*Oryx dammah*), maned wolf (*Chrysocyon brachyurus*), great apes (*Pan troglodytes, Gorilla gorilla, Pongo abelii*), gelada (*Theropithecus gelada*), American black bears (*Ursus americanus*) and Eurasian brown bears (*Ursus arctos*), and Eurasian beaver (*Castor fiber*) (14,31,37–40). This device collects continuous data on individual mean heart rate, physical activity, and body temperature on 2-minute, 15-minute, and 4-hour time scales, respectively, over spans of several months or even years (14,31,37–39). Baboons are a useful species for measuring real-time changes in heart rate in response to the environment, both because they are a common model for human cardiac physiology, and because considerable prior work links baboons’ social conditions to stress and health (41–43). In the wild, baboons live in multi-male, multi-female social groups, where individuals form strong, differentiated social relationships and access to resources is governed by strong “despotic” dominance hierarchies (43,44). Previous studies have demonstrated that baboons display physiological signals associated with chronic social and ecological stressors through fecal glucocorticoid measurements, but cardiac responses to these stressors are largely unknown (45–48).

Our specific objectives for this study were three-fold. First, we evaluated the safety, data collection duration, and accuracy of the Reveal LINQ^TM^ ICM in 10 baboons (*Papio anubis* and *P. cynocephalus*) at two implant locations on the body (see Methods). Second, we used the resulting data from the Reveal LINQ^TM^ ICMs to measure mean and resting heart rate, physical activity, and body temperature in freely moving captive baboons during their normal activities. We believe these are the first such data from unsedated or unrestrained baboons. Third, to test the biological validity of the ICM’s data, we assessed the extent to which two variables recorded by the ICM—heart rate and physical activity—were predicted by several internal and external variables, including body mass, time of day, ambient temperature, social dominance rank, and ovarian cycle phase.

We expected to observe the highest heart rates in the smallest individuals because of the well-known relationships between body size, heart rate, and metabolic rate (49,50). We also expected that baboon heart rate would show a circadian rhythm with the highest heart rates occurring during the day when baboons were physically active and ambient temperatures were warm, and lowest heart rates occurring at night when the animals are sleeping and temperatures are cool (51). We did not have strong expectations for how social dominance rank and ovarian cycle phase would predict heart rate, but low rank is linked to greater social stress in female baboons (43,44,48), and ovarian cycle phase is linked to changes in heart rate in humans (52,53). In terms of physical activity, we predicted that the baboons would be most active during the day, but we did not have strong expectations about how variation in activity would be correlated with ambient temperature, body mass, dominance rank, or ovarian cycle phase. Our results represent an important step to validate the use of the Reveal LINQ^TM^ ICM in baboons and to understand individual variation in physiology.

## METHODS

### Study subjects

Our study was conducted over an 8-month period, from January to September 2023. The subjects were 10 female baboons belonging to two closely related species: *Papio anubis* (i.e., olive baboons; n=5) and *P. cynocephalus* (yellow baboons; n=5). The animals were group-housed in species-appropriate enclosures to promote natural social behaviors at the Kenya Institute of Primate Research (KIPRE; each enclosure housed the five females of each species with no other residents). All females were identified using colored collars, individualized patterns of hair shaving, and unique physical traits, allowing us to distinguish individuals at all times regardless of their posture or location in the enclosure. All females were sexually mature adults that were caught in the wild at sites in Kenya by KIPRE staff; hence their ages were unknown. All females experienced natural ovarian cycles throughout the study, but because they were isolated from adult males, none became pregnant or had offspring. Animal care attendants visited each group daily to feed and clean the enclosures between 09:00 and 10:00. All animals were fed the same diet of monkey cubes (Unga Farm Care, Ltd., Nairobi, Kenya) and received regular supplementation of fresh fruits and vegetables (typically in the afternoon), and water was provided *ad libitum*. Veterinarians and veterinary technicians performed daily visual health evaluations and ovarian cycle phase checks. Animals did not change location during the study, with the exception of one individual (PCY216) who, for 5 days, was housed in a side enclosure adjacent to her groupmates’ enclosure while she was recovering from wound repair (data for PCY216 from this 5-day period are excluded from our analyses).

### Insertable cardiac monitor placement

All individuals received a Reveal LINQ^TM^ ICM (Medtronic, Minneapolis, MN, USA; 4.0 mm × 7.2 mm x 44.8 mm; mass 2.4 g), which was chosen due to its demonstrated safety and accuracy in other wild and captive mammalian species (14,31,37,38). Each ICM was loaded with the most recent version of “B-Ware” software, which recorded three types of data: (1) average heart rate in beats per minute (bpm) every 2 minutes; (2) total physical activity every 15 min (physical activity data is collected by an accelerometer in the ICM and is represented by the ICM as the number of “active minutes” in a 15-minute period); and (3) average body temperature in Celsius every 4 hours (39). Additionally, the ICM recorded an ECG strip for all tachycardia and bradycardia episodes the baboon experienced as defined by thresholds set during implant programming (see below).

Our implantation procedure was similar to previously published protocols (14,31,37,38). Baboons were sedated using a mixture of ketamine hydrochloride (HCl) and xylazine at a dose of 10 mg/kg ketamine-HCl and 0.5 mg/ml xylazine administered intramuscularly (adding 0.5 ml of 100 mg/ml xylazine to a 10-ml bottle of ketamine [100 mg/ml], dosed at 0.1 ml/kg; dose was estimated from the animal’s body mass at their most recent previous sedation). After sedation, we measured body mass (to the nearest 0.1 kg), rectal body temperature, respiration rate (respiration rate was measured as breaths per minute over 1 minute and was collected solely for monitoring purposes), and crown-rump length (the length of the baboon from the top of the head to the base of the tail measured while the baboon was supine). All animals were closely monitored during anesthesia by the project veterinarians and technicians, with vital signs assessed before, during, and after the implant procedure to ensure their safety, well-being, and proper healing of the implant site.

Prior to implantation, each animal was transported to a surgical preparation room and positioned to expose the left side of their chest. The animal was then shaved at one of two body sites that we considered for candidate implant locations: the left pectoral muscle between ribs 2 and 4 (“pectoral placement”), and the lower chest over the left side ribs, between ribs 5 and 8 (“lateral placement”; **Figure 1**). At each site, we assessed electrocardiogram (ECG) R-wave amplitude using real time recordings from a KardiaMobile 6L^TM^ personal ECG machine (Kardia© by AliveCor Inc., Mountain View, CA, USA). Based on the identification of sites with optimal R-wave amplitude, we marked an incision site parallel to the edge of the Kardia on the animal’s chest. Afterwards, the animals were transferred to a sterile surgical suite where six baboons received pectoral implants, and four baboons received lateral implants. Each animal was assigned an implant location based on the strength their R-wave amplitudes at each site and to attain roughly equal number of females with an ICM at either location between both species.

**Figure 1.**
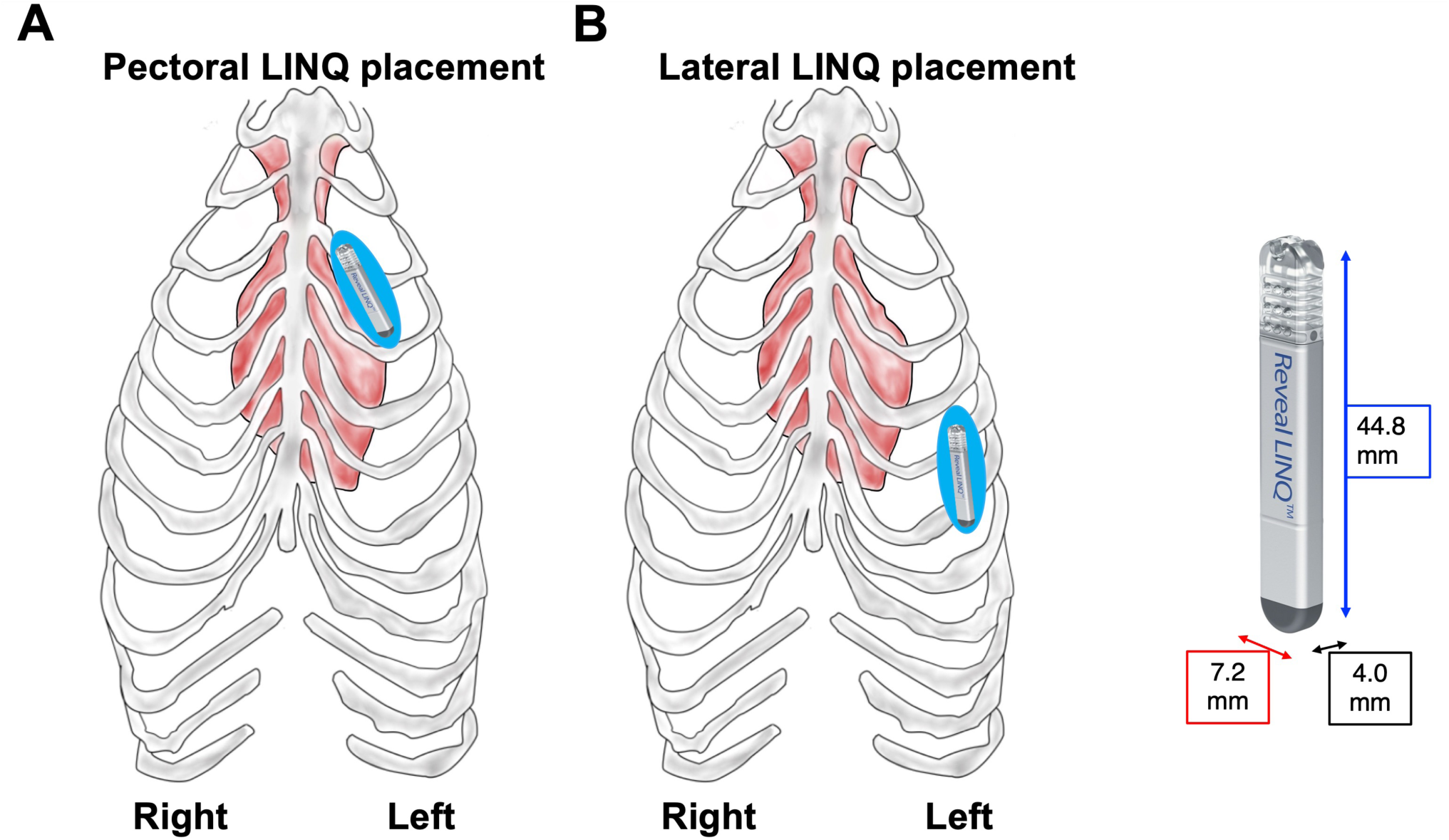
**Locations for subcutaneous Reveal LINQ**^TM^ **placement.** Illustration (**A**) shows the position of pectoral LINQ placement over the intercostal spaces between ribs 2 through 4. Illustration (**B**) shows the position of lateral LINQ placement over the intercostal spaces between ribs 5 through 8. In both (**A**) and (**B**), right and left rib labels are included to help define the perspective taken in the illustration. This illustration is intended to show device placement only and is not drawn to scale. An image of the Reveal LINQ^TM^ is shown to the right along with its dimensions (44.8 mm x 7.2 mm x 4.0mm). Each dimension is denoted by a different color. Device length (44.8 mm) is in blue, width (7.2 mm) is denoted in red, and depth (4.0mm) is in black.

After selecting the implant location, we cleaned and prepared the site with Betadine antiseptic solution (10% povidone-iodine). We then marked the planned incision site on the chest of the baboon with non-toxic marker and made a 0.5 cm incision perpendicular to the planned insertion site using Medtronic’s provided pre-sterilized insertion tools. Next, we inserted the provided blue “insertion tool” parallel to the body to create a ∼10 cm subcutaneous “pocket” into which we implanted the Reveal LINQ^TM^. The incision was closed with a simple interrupted suture pattern (using VeterSut VetCRYL Absorbable Polyglactin Surgical Suture USP Size 2-0, (SH) ½ 26mm Taper Point Needle, by VeterSut, Fort Myers, FL, USA) and Betadine antiseptic ointment (10% povidone-iodine) was applied to the site. Each animal was given antibiotic via intramuscular injection (procaine penicillin G 100,000 iu/ml, benzalline penicillin G 100,000 iu/ml dihydrostreptomycin sulphate 200mg/ml, 1cc/10kg body mass).

Following surgery, we programmed the animal’s ICM using a Medtronic 2090 Programmer and telemetry head. We began by synchronizing the internal clock of the ICM to the local time based on the time reported by the National Institute of Standards and Technology (NIST; https://timegov.nist.gov/). We then set thresholds for the ICM to identify tachycardia and bradycardia episodes based on parameters used for Reveal LINQ^TM^ ICMs in Gelada baboons (*Theropithecus gelada*) (38). Tachycardia episodes were defined as heart rate events ≥130 beats per minute (bpm) for at least 12 heart beats. Bradycardia episodes were defined as heart rate events of ≤50 bpm for at least 4 heart beats. Finally, we paired each Reveal LINQ^TM^ implant to a MyCareLink Patient Monitor^TM^ (Medtronic, Minneapolis, MN, USA), which was used to access data on tachycardia and bradycardia episodes (see below).

### Data downloads

To access data collected by the Reveal LINQ^TM^ ICM, we carried out monthly data downloads via direct telemetry using two devices: the Medtronic 2090 programmer and the MyCareLink Patient Monitor^TM^ (**Figure S1**). Because direct telemetry downloads required physical contact between the telemetry head and the skin over the healed implant site, baboons were sedated using a mixture of ketamine hydrochloride (10.0 mg/kg) and xylazine (0.5 mg/kg). Baboons were also weighed (to the nearest 0.1kg) during each download. As shown in **Figure S1**, the Medtronic 2090 programmer was used to download data recorded via B-ware: 2-minute mean heart rate (bpm), the number of active minutes in a 15-minute period, and 4-hour mean body temperature (°C). The MyCareLink Patient Monitor^TM^ was used to download ECG strips of tachycardia and bradycardia episodes. The ECG strip for each episodes contains beats that occur before and after the episode, a mark for the detected episode start beat, and a mark for the detected episode end beat. At each download, we also recorded both the ICM’s reported time and the true time at our location as reported by the NIST (the ICM’s time drifted relative to true time; see below for how we addressed this drift).

### Collecting data on predictors of heart rate and activity

To test how several variables predicted individual variation in heart rate and physical activity, we also required information on the animals’ environments and physical states. Below we describe how data on these variables were collected.

*Baboon species* was whether the female was *P. anubis* or *P. cynocephalus*.

*Implant location* was defined as the anatomical location where the Reveal LINQ^TM^ implant was inserted, either “pectoral” or “lateral” (**Figure 1**).

*Body mass* was calculated for each animal following as the animal’s weight in kilograms (kg), measured at each data download. Body mass measurements for each individual were time matched to heart rate and activity data points by date.

To model circadian rhythm, we defined *daytime* as the hours between median sunrise and median sunset times recorded throughout the study, 06:35 to 18:41. *Nighttime* was defined as the time when natural illumination from the sun was absent due to sunset (the time after sunset and before sunrise) between 18:41 to 06:35. Times of sunrise and sunset for the dates of the study were acquired for the location of KIPRE from the NOAA Solar Calculator (https://gml.noaa.gov/grad/solcalc/).

*Ambient temperature* was collected via a Kestrel DROP D2 Wireless Temperature & Humidity Data Logger (Kestrel Instruments, Nielsen-Kellerman Company, Boothwyn, PA, USA) programmed to log temperature every 30 minutes at the animal enclosures.

*Dominance rank* was determined by assigning wins and losses in dyadic agonistic interactions collected during 20-minute focal animal behavioral observations conducted by KIPRE ethology staff. Each of the ten females were sampled several times per week over the 8-month study period (251 total hours of observation across all 10 females). We entered agonistic interactions into a dominance win-loss matrix including the five females in their enclosure (i.e., the five females in the *P. cynocephalus* enclosure were ranked separately from the five females in the *P. anubis* enclosure). In each matrix, individuals were placed in order of descending rank to minimize the number of entries that fall below the diagonals of the matrices. An individual with a rank of “1” is most dominant and an individual with a rank of “5” is the least dominant in her group.

*Ovarian cycle phase* was determined from the KIPRE animal care staff’s daily visual assessment of the baboons. Adult female baboons experience a 30–35-day ovarian cycle, which contains four distinct phases (follicular, periovulatory, luteal, menstruation) (54,55). These phases were determined using well-validated changes in sexual skin size and visual evidence of menstruation (56).

### Data processing

Before starting our analyses, we processed the ICM data in two ways. First, we adjusted for offsets in the ICM’s recorded time. During each monthly data download we noted that the time reported by the Reveal LINQ^TM^ differed from the real time by several seconds. This drift created a time offset such that each ICM’s recorded time differed from the true local time (average monthly offset = 132 seconds; range = 9 to 938 seconds). We assumed a constant rate of drift. For example, if the data collected by the ICM were off by 30 seconds on the first download day, and 90 seconds on the second download day 30 days later, then the daily offset rate between downloads would be 2 seconds (60 seconds divided by 30 days = 2 seconds). We would then assume that the data collected on day 2 by the ICM would be offset by 32 seconds, on day 3 by 34 seconds and so on. We used this daily offset value to adjust the time obtained from the ICMs to match the inferred local time.

Second, we excluded any data that was recorded on a day that a baboon underwent a sedated medical procedure and the 5-day period when one individual was housed in a side enclosure (see above). We did this to control for the potential effects of sedative drugs on individual physiology as well as the potential effects of psychosocial stress that an individual may experience while undergoing a change in their social and physical environment.

### Statistical analyses

#### Objective 1

To test the ICM’s ability to accurately sense heart rate, we used two main approaches based on those used in prior studies (14,23). First, we noted the amplitudes (in millivolts, mV) of the ICM’s R-, T- and P-waves from 100 randomly selected ECG strips collected during tachycardia and bradycardia episodes. We then compared the amplitudes of the R-waves to the T- and P- waves that co-occurred on an ECG strip (10 ECG strips per female; **Figure S2**). This test is important because especially strong T- and P-waves can create interference that hinders the ICM’s ability to measure R-wave intervals and calculate heart rate. To test whether ECG R-, P-, and T- wave amplitudes differed as a function of implant site (pectoral or lateral) and baboon species (*P. anubis* or *P. cynocephalus*), we also ran a series of Welch’s t-tests for each category of ECG wave.

Second, to test whether the heart rates reported by the ICM were accurate, we correlated the R-wave count reported by the ICM on the same 100 ECG strips described above with manually determined (hand-counted) R-waves from the same strips (10 ECG strips per baboon). We then correlated the ICM’s reported R-wave counts with the manual R-wave counts using a Pearson’s correlation using the R package *corr* (57). To test whether sensing accuracy varied as a function of implant site (pectoral or lateral) and baboon species (*P. anubis* and *P. cynocephalus*), we ran a two-way ANOVA using the aov() function in R.

#### Objective 2

To provide baseline data on captive baboon cardiac physiology, we identified the maximum and minimum observed heart rates recorded during all tachycardia and bradycardia episodes detected by the Reveal LINQ^TM^ for all subjects. Specifically, we identified each clearly defined R-wave peak from all tachycardia/bradycardia ECG strips collected from each individual over the study period. We then used the R-R interval reported in milliseconds (ms) between each set of consecutive discernible ECG waveforms (**Figure S2**) to calculate the maximum observed heart rate in bpm for the tachycardia/bradycardia episode using the following formula: 60000/R-R interval (ms) = heart rate (bpm). In all subsequent analyses, we excluded any 2-minute mean heart rate values that were higher than the observed maximum heart rate and any 2-minute heart rate values that were lower than the observed minimum heart rate.

We also calculated each female’s mean daytime, nighttime, and resting heart rate from the data on 2-minute mean heart rates over the study period for each baboon. A given 2-minute mean heart rate value was assigned to daytime or nighttime using the definitions described above. Average daily resting heart was calculated by averaging the 2-minute mean heart rate recordings between 05:30 and 05:45. Because the 2-minute period of the heart rate measures did not neatly overlap the 15-minute period, we weighted the starting bin of the 2-minute heart rate recording one minute outside of this period by.25 and the ending bin one minute after this period by.25. We did this to account for the variation in our data as to which heart rate measurement contributed the additional “1-minute” of heart rate data for the 15-minute period corresponding to the activity data point. We chose 05:30 and 05:45 to measure resting heart rate because it corresponded to the lowest nighttime heart rate readings before the animals climbed down from their sleeping perches and began being active for the day (**Figure S3**).

We also calculated each female’s average daytime and nighttime activity levels as the mean number of active minutes in a 15-minute period during day or night (following the definitions above). Finally, we report each female’s average 4-hour body temperature over the course of the study.

#### Objective 3

To test how heart rate and physical activity were predicted by external and internal variables, we ran a series of linear mixed models (LMMs) in R using the packages lme4 (58) and lmerTest (59). For models of heart rate, we aggregated the 2-minute mean heart rate data to a 15-minute scales to allow us to control for physical activity, which was measured by the Reveal LINQ^TM^ on a 15-minute scale (activity is an essential covariate for understanding variation in heart rate). The activity metric is reported as the number of “active minutes” in a 15-minute period and was calculated by the Reveal LINQ^TM^ using a proprietary algorithm. Our response variable was 15-minute mean heart rate, calculated as the average of the 2-minute mean heart rates surrounding this 15-minute period and weighting the starting and ending bins using the same approach described above for resting heart rate. The fixed effects in these models were the number of active minutes in a 15-minute period, baboon species, implant location, body mass, whether a data point was collected in the daytime or nighttime, mean ambient temperature (°C) during the 30 minutes when the data were collected, dominance rank over the 9-month course of the study (ranks did not change over time), and ovarian cycle phase on the day of the observation. We modeled baboon identity as our random effect. We identified the best-fitting model using Akaike Information Criterion (AIC) implemented in the aictab() function in the AICcmodavg package (60). The best-fitting model was >2 AIC units from all possible models.

Likewise, we ran a series of LMMs to test how activity levels were predicted by a range of external and internal variables. Our response variable was the number of active minutes in a 15-minute period. The fixed effects were baboon species, implant location, body mass, whether a data point was collected in the daytime or nighttime, ambient temperature (°C) during the 30-minute period when the data were collected, dominance rank over the 9-month course of the study (ranks did not change over time), and ovarian cycle phase on the day of observation. Baboon identity was included as a random effect. As above, the best-fitting model was >2 AIC units from all possible models.

## RESULTS

### Objective 1: ICM safety, data duration, and accuracy

**Table 1** describes the 10 animals who received Reveal LINQ^TM^ ICM implants, their implant locations (pectoral or lateral), and their duration of data collection. Following implantation, none of the animals were observed to interfere with the ICM implant site, none of the ICM implant sites showed signs of infection, and all animals healed without complications. However, one ICM was rejected by one animal 33 days after implantation (*P. cynocephalus* individual PCY212). This animal exhibited no signs of infection prior to rejection (e.g. no redness or swelling at the incision site; no rise in body temperature recorded by the ICM). The ICM was recovered from the floor of the animal’s enclosure, sterilized, and surgically reimplanted 23 days later in the same location as the animal’s first implantation (the lateral location). No signs of infection or complications occurred after re-implantation, and the ICM continued to collect data from PCY212 until the end of its battery life.

**Table 1.**
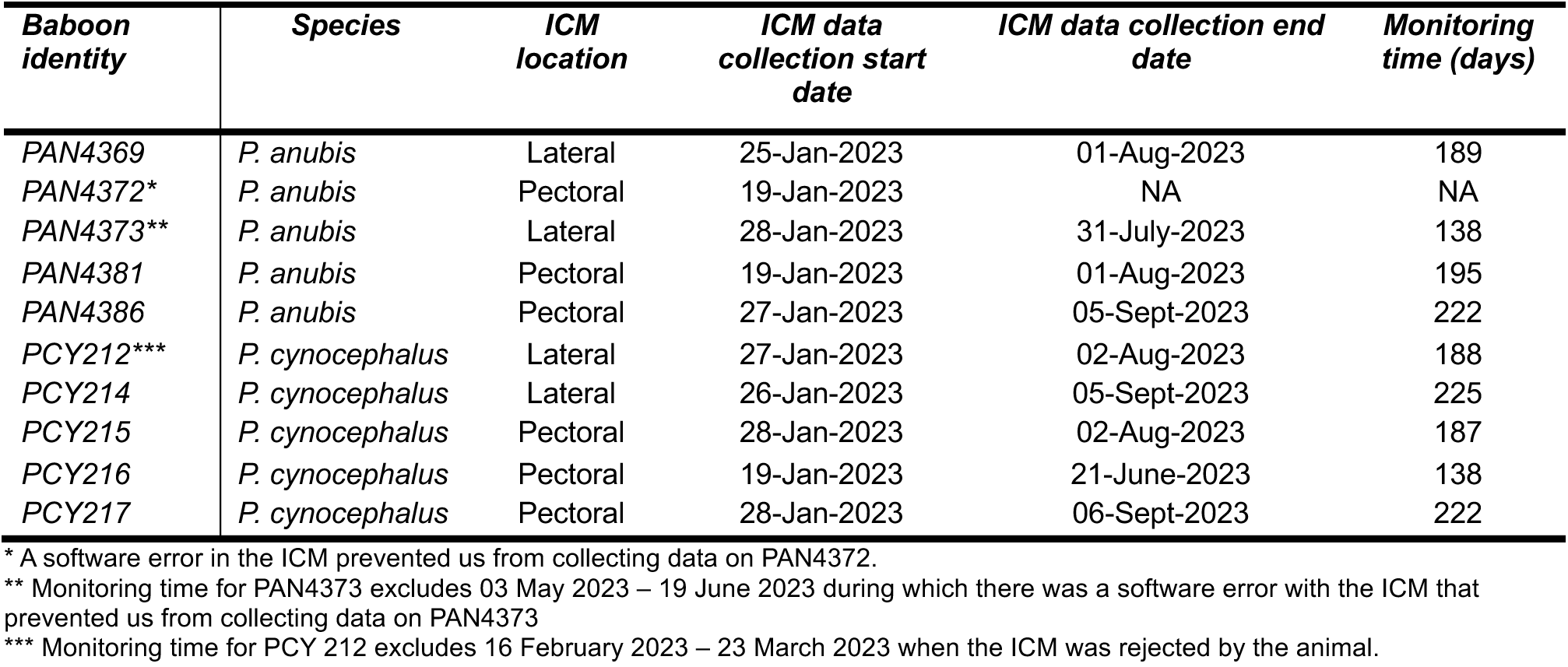
Study subjects and ICM data collection duration, including baboon species, anatomical location of their ICM, the date their ICM was inserted, the date their ICM’s battery died, and the duration of ICM data collection in days.

| Baboon identity | Species | ICM location | ICM data collection start date | ICM data collection end date | Monitoring time (days) |
| --- | --- | --- | --- | --- | --- |
| PAN4369 | *P. anubis* | Lateral | 25-Jan-2023 | 01-Aug-2023 | 189 |
| PAN4372* | *P. anubis* | Pectoral | 19-Jan-2023 | NA | NA |
| PAN4373** | *P. anubis* | Lateral | 28-Jan-2023 | 31-July-2023 | 138 |
| PAN4381 | *P. anubis* | Pectoral | 19-Jan-2023 | 01-Aug-2023 | 195 |
| PAN4386 | *P. anubis* | Pectoral | 27-Jan-2023 | 05-Sept-2023 | 222 |
| PCY212*** | *P. cynocephalus* | Lateral | 27-Jan-2023 | 02-Aug-2023 | 188 |
| PCY214 | *P. cynocephalus* | Lateral | 26-Jan-2023 | 05-Sept-2023 | 225 |
| PCY215 | *P. cynocephalus* | Pectoral | 28-Jan-2023 | 02-Aug-2023 | 187 |
| PCY216 | *P. cynocephalus* | Pectoral | 19-Jan-2023 | 21-June-2023 | 138 |
| PCY217 | *P. cynocephalus* | Pectoral | 28-Jan-2023 | 06-Sept-2023 | 222 |

\* A software error in the ICM prevented us from collecting data on PAN4372.
\*\* Monitoring time for PAN4373 excludes 03 May 2023 – 19 June 2023 during which there was a software error with the ICM that prevented us from collecting data on PAN4373
\*\*\* Monitoring time for PCY 212 excludes 16 February 2023 – 23 March 2023 when the ICM was rejected by the animal.

On average, the ICMs collected data for 191 days (range=138 to 225 days; **Table 1**). We were unable to download data from one ICM (*P. anubis* individual PAN4372) due to a problem with the ICM’s software that we were unable to resolve; hence all subsequent analyses use data from 9 individuals. For these 9 devices, data collection concluded when the batteries died (**Table 1**).

The ECGs reported by the ICM had strong R-wave amplitudes: the mean R-wave amplitude across all subjects was 0.29 mV, which was 6.3 times the amplitude of the T-wave, and 3.1 times the amplitude of the P-wave (mean T-wave amplitude=0.05 mV; mean P-wave amplitude=0.11 mV; N=100 ECGs; 10 each from 10 baboons; **Table 2**). ECG wave amplitudes varied as a function of baboon species such that *P. cynocephalus* individuals displayed larger P-and T-wave amplitudes, and perhaps larger R-wave amplitudes compared to *P. anubis* individuals (**Table S1:** T-wave: t=-2.08, p=0.041; P-wave: t=-2.77, p=0.007; R-wave: t=-1.99, p=0.051). ECG wave amplitudes did not vary as a function of ICM implant site (R-wave: t=0.043, p=0.966; T-wave: t=1.21, p=0.232; P-wave: t=-0.156, p=0.879; **Table S1**).

**Table 2.** Heart rate and Reveal LINQ^TM^ ECG amplitude metrics for 9 female baboons, including the baboon’s species, mean daytime heart rate (HR), mean nighttime heart rate, mean resting heart rate, and mean R, P, and T-wave amplitudes. We also report the Pearson’s correlation between the Reveal LINQ^TM^ reported HR with manually determined (hand-counted) HR.

| Baboon identity | Species | Mean R-wave amplitude (mV) | Mean P-wave amplitude (mV) | Mean T-wave amplitude (mV) | Pearson's *r* for heart rate validation | Resting HR (bpm, ± SD) | Mean daytime HR (bpm, ± SD) | Mean nighttime HR (bpm, ± SD) | Mean daytime activity (active minutes/15-min period, ± SD) | Mean nighttime activity (active minutes/15-minute period, ± SD) | Mean body temperature ($^{\circ}$C, ± SD) |
| --- | --- | --- | --- | --- | --- | --- | --- | --- | --- | --- | --- |
| PAN4369 | *P.anubis* | 0.200 | 0.023 | 0.051 | 0.995 | 76.07<br>(9.03) | 94.00<br>(15.90) | 81.30<br>(14.80) | 3.73<br>(3.73) | 0.38<br>(1.32) | 36.80<br>(1.70) |
| PAN4372* | *P.anubis* | 0.315 | 0.045 | 0.093 | 0.999 | NA | NA | NA | NA | NA | NA |
| PAN4373** | *P.anubis* | 0.235 | 0.035 | 0.083 | 0.998 | 77.04<br>(11.39) | 96.30<br>(21.40) | 84.00<br>(16.80) | 4.48<br>(3.86) | 0.35<br>(1.16) | 36.70<br>(0.52) |
| PAN4381 | *P.anubis* | 0.290 | 0.043 | 0.090 | 0.999 | 98.23<br>(8.53) | 122.0<br>(17.4) | 107.0<br>(11.10) | 4.58<br>(3.65) | 0.33<br>(1.09) | 35.9<br>(0.70) |
| PAN4386 | *P.anubis* | 0.330 | 0.053 | 0.075 | 0.996 | 76.61<br>(7.47) | 89.70<br>(16.50) | 78.60<br>(12.20) | 3.09<br>(2.54) | 0.22<br>(0.88) | 36.20<br>(0.60) |
| PCY212*** | *P.cynocephalus* | 0.295 | 0.050 | 0.100 | 0.999 | 94.57<br>(7.54) | 111.00<br>(18.00) | 103.00<br>(12.40) | 2.18<br>(2.25) | 0.20<br>(0.81) | 36.50<br>(0.79) |
| PCY214 | *P.cynocephalus* | 0.365 | 0.055 | 0.100 | 0.944 | 82.10<br>(8.91) | 107.00<br>(18.70) | 87.70<br>(15.80) | 1.15<br>(2.04) | 0.18<br>(0.92) | 36.70<br>(1.80) |
| PCY215 | *P.cynocephalus* | 0.300 | 0.055 | 0.105 | 0.999 | 93.94<br>(9.98) | 128.00<br>(19.80) | 109.00<br>(19.30) | 1.99<br>(2.33) | 0.21<br>(0.88) | 36.30<br>(0.70) |
| PCY216 | *P.cynocephalus* | 0.260 | 0.058 | 0.095 | 0.995 | 96.72<br>(9.14) | 118.00<br>(17.90) | 106.00<br>(13.90) | 2.35<br>(2.51) | 0.16<br>(0.62) | 36.50<br>(0.90) |
| PCY217 | *P.cynocephalus* | 0.325 | 0.073 | 0.150 | 0.993 | 74.36<br>(6.17) | 99.60<br>(23.80) | 84.20<br>(17.00) | 2.28<br>(2.16) | 0.29<br>(0.94) | 36.30<br>(0.64) |

\* A software error in the ICM prevented us from collecting data on PAN4372.
\*\* Monitoring time for PAN4373 excludes 03 May 2023 – 19 June 2023 during which there was a software error with the ICM that prevented us from collecting data on PAN4373
\*\*\* Monitoring time for PCY 212 excludes 16 February 2023 – 23 March 2023 when the ICM was rejected by the animal.

The R-wave counts recorded by the Reveal LINQ^TM^ were highly correlated with R-wave peaks we counted by hand from representative ECGs (**Figure 2**). The Pearson’s R between ICM-reported and manual (hand-counted) peaks ranged from 0.999 to 0.944 across all individuals (**Table 2**). Sensing accuracy did not vary by species (ANOVA: difference between *P. anubis* and *P. cynocephalus* =-1.24 bpm, p=0.34). However, sensing accuracy differed by implantation site: the three worst performing ICMs included 3 of the 4 laterally placed ICMs (ANOVA: difference between pectoral and lateral sensing=-5.967 bpm, p<0.001; **Table 2**; **Figure 2A**). Laterally placed ICMs reported 6 fewer beats than the manually counted (hand-counted) beats when compared to pectorally placed implants.

**Figure 2.**
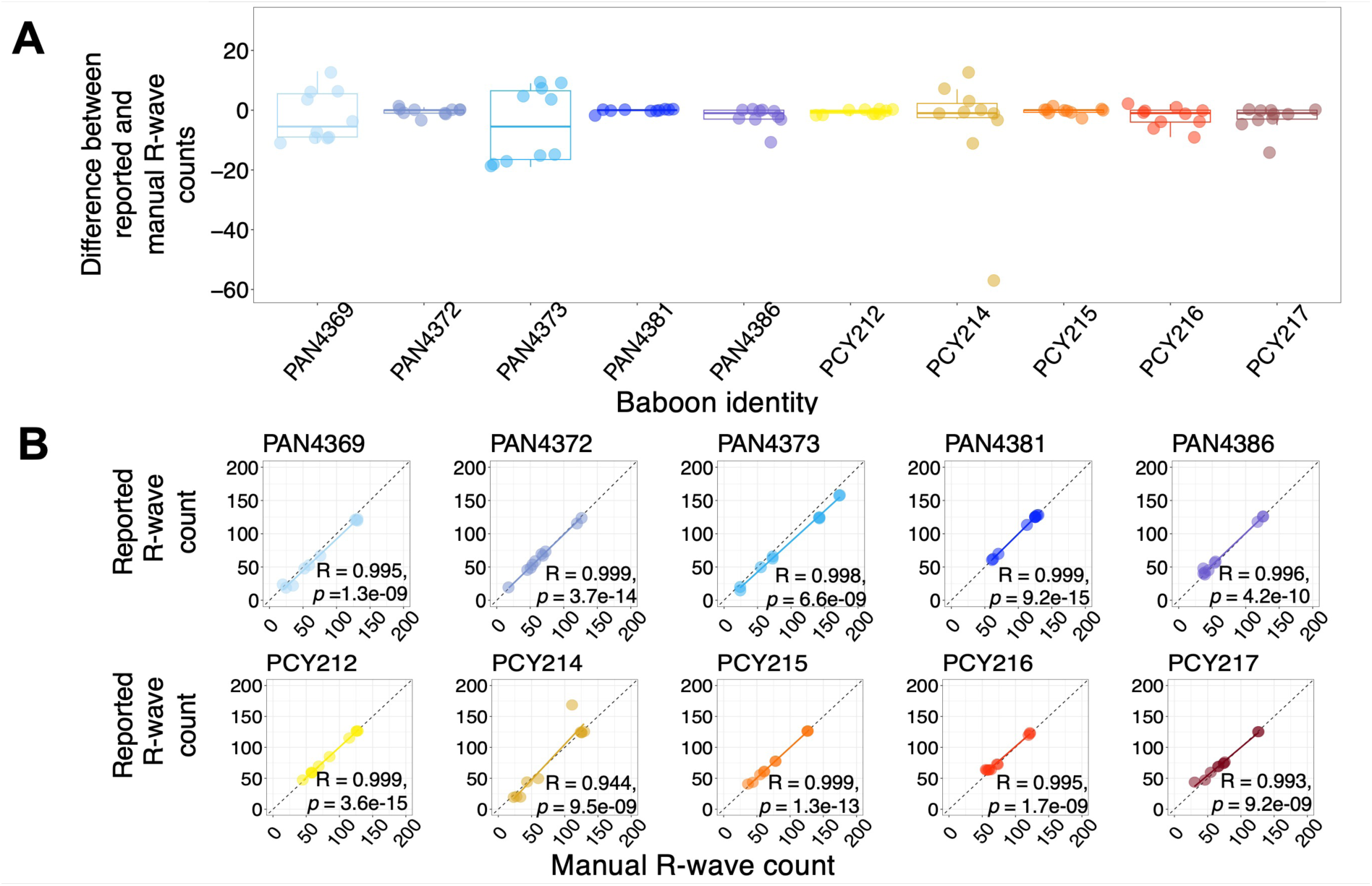
**Reveal LINQ**^TM^ **ICMs accurately detect heart rate in baboons.** Panel (**A**) shows the difference between the ICM’s reported and manual R-wave counts (y-axis) for 10 ECG strips from each subject (x-axis). Each point represents the difference in these R-wave counts for 1 ECG strip; positive numbers represent higher ICM-reported than manually counted peaks; negative numbers represent higher manually counted than ICM-reported peak counts. Each individual baboon has a distinct color and color gradients represent baboon species; blue to purple points and lines represent *P. anubis* and yellow to red points and lines represent *P. cynocephalus.* The middle line of the boxplot represents the median value of difference between reported and manual R-wave peak counts. The bottom and top whiskers represent the minimum and maximum difference in these values respectively. The bottom and top edges of the box represent the 25^th^ and 75^th^ percentiles. Panel (**B**) shows the correlation between reported and manual peak counts for 10 ECG strips per baboon. Baboon identity is listed above each plot.

Pearson’s R ranged from 0.999 to 0.944 across subjects. The dashed line represents the 1:1 line. As in panel A, color represents baboon identity and species.

### Objective 2: Physiological parameters for captive female baboons

**Figure 3** shows representative data on 2-minute mean heart rate and the number of active minutes in a 15-minute period recorded by ICMs over a one-week period from two baboons (one *P. cynocephalus* (PCY215) and one *P. anubis* (PAN4386).

**Figure 3.**
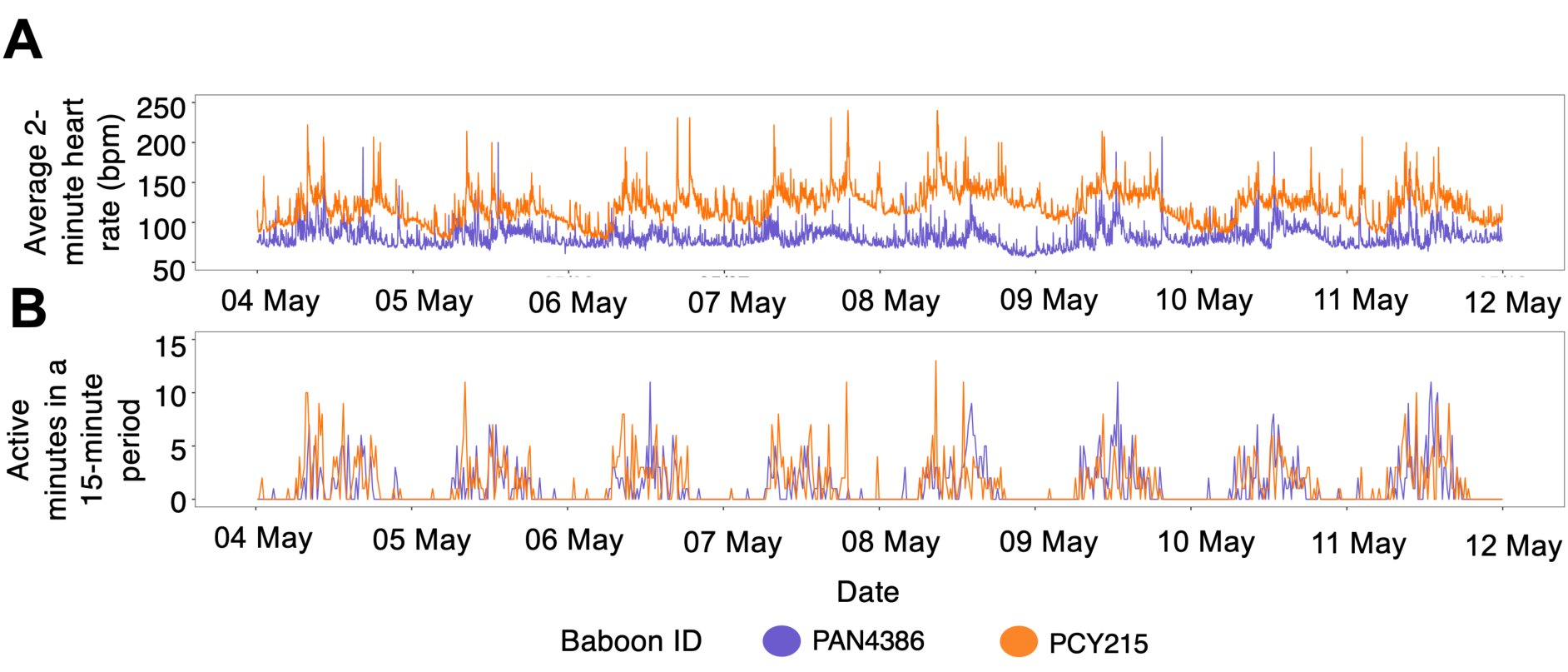
Representative data on 2-minute mean heart rate and 15-minute activity levels for two baboons. Panel (**A**) shows longitudinal changes in 2-minute mean heart rate (bpm) captured by the Reveal LINQ^TM^ for two females between 04 May 2023 and 11 May 2023. Purple lines show data from individual PAN4386 (*P. anubis*) and orange lines show data from individual PCY215 (*P. cynocephalus*). Panel (**B**) shows longitudinal change in the number of active minutes recorded in a 15-minute period for the same two females over the same period of time as panel A.

The maximum observed HR occurred within a 12-beart tachycardia episode for PCY214 where the mean HR across all 12 beats was 228.6 bpm (mean=262.5 ms between consecutive R-waves; **Figure S4**). The maximum observed HR detected between two consecutive R-waves in this episode was 260 bpm (230 ms between R-waves; **Figure S4**). This tachycardia event occurred while PCY214 and her groupmates were being sedated for their monthly data download (note that data during sedations were excluded from our other analyses as described in the methods). The minimum observed HR occurred within a 4-beat bradycardia episode experienced by PAN4372 where the mean HR across all 4 beats was 48.1 bpm (mean=1245 ms between consecutive R-waves; **Figure S5**), and the minimum interval between two consecutive R-waves was 47.2 bpm (1270 ms between R-waves; **Figure S5**). We note that a slightly slower R-R interval was observed prior to the Reveal LINQ’s detection of the 4-beat bradycardia episode (46.5 bpm or 1290 ms between two consecutive R-waves; **Figure S5**). Contextually, the ECG strip that includes this bradycardia event and the HR values that precede and follow it was recorded during the nighttime hours (04:35) when PAN4372 was likely sleeping.

For each individual baboon, the maximum observed heart rate for a single R-R wave interval ranged from 139.5 to 260 bpm in tachycardia episodes and the minimum observed heart rate ranged from 47.2 to 49.6 bpm in bradycardia episodes. Average daily resting heart rate (measured between 05:30 and 05:45), was highly variable between baboons, ranging from 74.36 bpm (SD±6.17 bpm in PCY217) to 98.23 bpm (SD±8.53 bpm in PAN4381; **Table 2**). Average individual daytime heart rate ranged from 89.70 bpm (SD±16.50 bpm) to 128.00 bpm (SD±19.80 bpm) while average individual nighttime heart rate ranged from 78.60 bpm (SD± 12.20 bpm) to 109.00 bpm (SD±19.30 bpm) (**Table 2**).

In addition to providing information on heart rate, we also gathered data on physical activity levels and body temperature for the nine female subjects. The average daily number of active minutes in a 15-minute period across all nine subjects was 1.56 minutes (SD±2.61 minutes). Baboons were more active during the day than at night: during the daytime, the average number of active minutes in a 15-minute period ranged from 1.15 minutes (SD±2.04 minutes) in the least active individual to 4.58 minutes (SD±3.65 minutes) in the most active individual, while at night, the average number of active minutes in a 15-minute period ranged from 0.16 minutes (SD±0.62 minutes) to 0.38 minutes (SD±1.32 minutes; **Table 2**). Average body temperature over the 8-month study period varied little across subjects (range=35.90°C to 36.80°C) and was, on average, 36.40°C (SD± 1.08 °C) across all nine subjects (**Table 2**).

### Objective 3: Predictors of heart rate in captive female baboons

We used an information theoretic approach to test which variables best explained variation in 15-minute mean heart rate (HR) across all nine baboons. The best-supported model included all fixed and random effects (ι1AIC >2 compared to all other models; **Table 3; Table S2**). One of the most important variables was female identity, which explained 41% of the variation in 15-minute mean HR. Controlling for this variation, HR was also predicted by baboon species, implant location, body mass, dominance rank, ovarian cycle phase, and the number of active minutes in the 15-minute period (**Table 3**). HR was also explained by ambient temperature (higher HR at higher ambient temperatures) and whether the HR was recorded during the daytime or nighttime (**Table 3**). As expected, heart rate was positively correlated with the number of active minutes in the 15-minute period (**Figure S6**). A visualization of circadian variation in HR showed that, the lowest heart rates occurred between 23:00 to 06:00, before rapidly increasing at around 06:00, coinciding with the start of daylight (**Figure 4A**). Heart rate peaked between 09:00 and 10:00 for all individuals and remained relatively high until 18:00 when dusk approached and heart rate began to decrease. We also visualized the effects of ambient temperature and individual body mass, which are shown in **Figures S7 and S8.**

**Figure 4.**
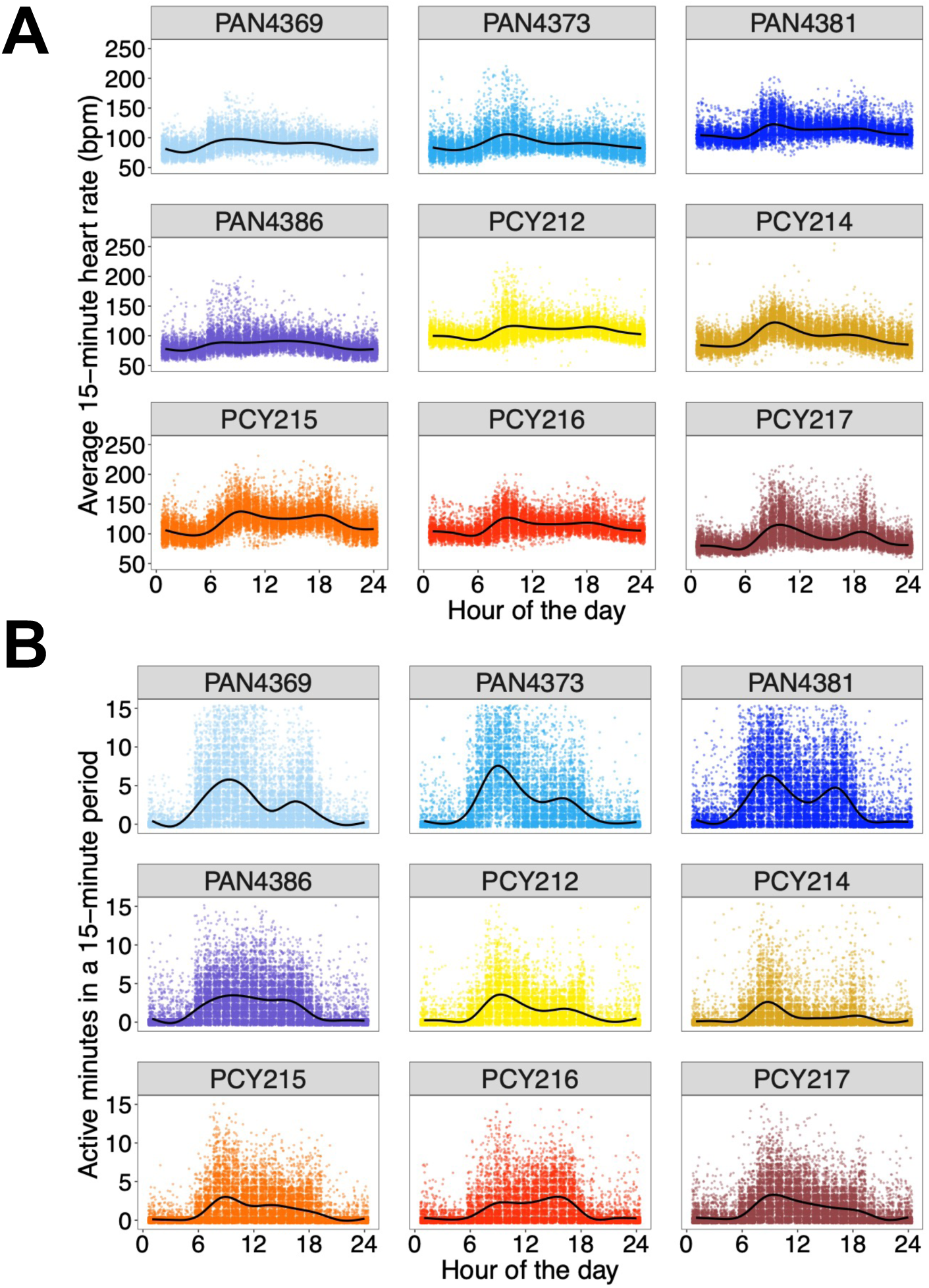
Baboons display circadian variation in heart rate and activity. Plots show the relationship between time of day (x-axis) and either (**A**) average 15-minute heart rate in bpm or (**B**) the number of active minutes in a 15-minute period. Each sub-plot shows these relationships for an individual baboon; female identity is at the top of each sub-plot. The sub-plots were made using data collected from each female’s ICM data collection start date to data collection end date, excluding days of sedated procedures and other anomalies as noted in Table 1. Colors represent individual identity and gradients represent baboon species; blue to purple points represent *P. anubis* and yellow to red points represent *P. cynocephalus*.

**Table 3.** Summary of the best-supported LMM (HR9 in **Table S2**) explaining variation in heart rate in female baboons at KIPRE. This model was >2 AIC units from all other models (**Table S2**). Reference categories for each fixed effect are denoted in parentheses.

| *Fixed effects* | *Estimate*<br>*(± SE)* | *T-value* |
| --- | --- | --- |
| *Species (P. cynocephalus)* | 7.41 (8.42) | 0.88 |
| *ICM implant location (pectoral)* | 9.21 (9.430) | 0.98 |
| *Body mass (kg)* | -1.07 (0.05) | -22.57 |
| *Day or night* | -10.49 (0.10) | -103.13 |
| *Ambient temperature (°C)* | 0.31 (0.01) | 29.91 |
| *Dominance rank* | -3.23 (3.13) | -1.03 |
| *Cycle phase: Periovulatory (Follicular)* | 0.25 (0.09) | 2.84 |
| *Cycle phase: Luteal (Follicular)* | 1.51 (0.13) | 11.53 |
| *Cycle phase: Menstruation (Follicular)* | -0.89 (0.14) | -6.07 |
| *Number of active minutes in 15-minute period* | 0.61 (0.02) | 34.96 |

Low-ranking individuals had lower average 15-minute heart rates than more dominant individuals such that a 1-step increase ordinal dominance rank (i.e., a lower rank) was linked to a 3-bpm reduction in 15-minute mean HR (**Figure 5A**; **Table 3**). Individual heart rate was higher when individuals were in the periovulatory and luteal phases of the ovarian cycle, and lower during menstruation, compared to the follicular phase, although this effect was small (**Figure 5B**; **Table 3**).

**Figure 5.**
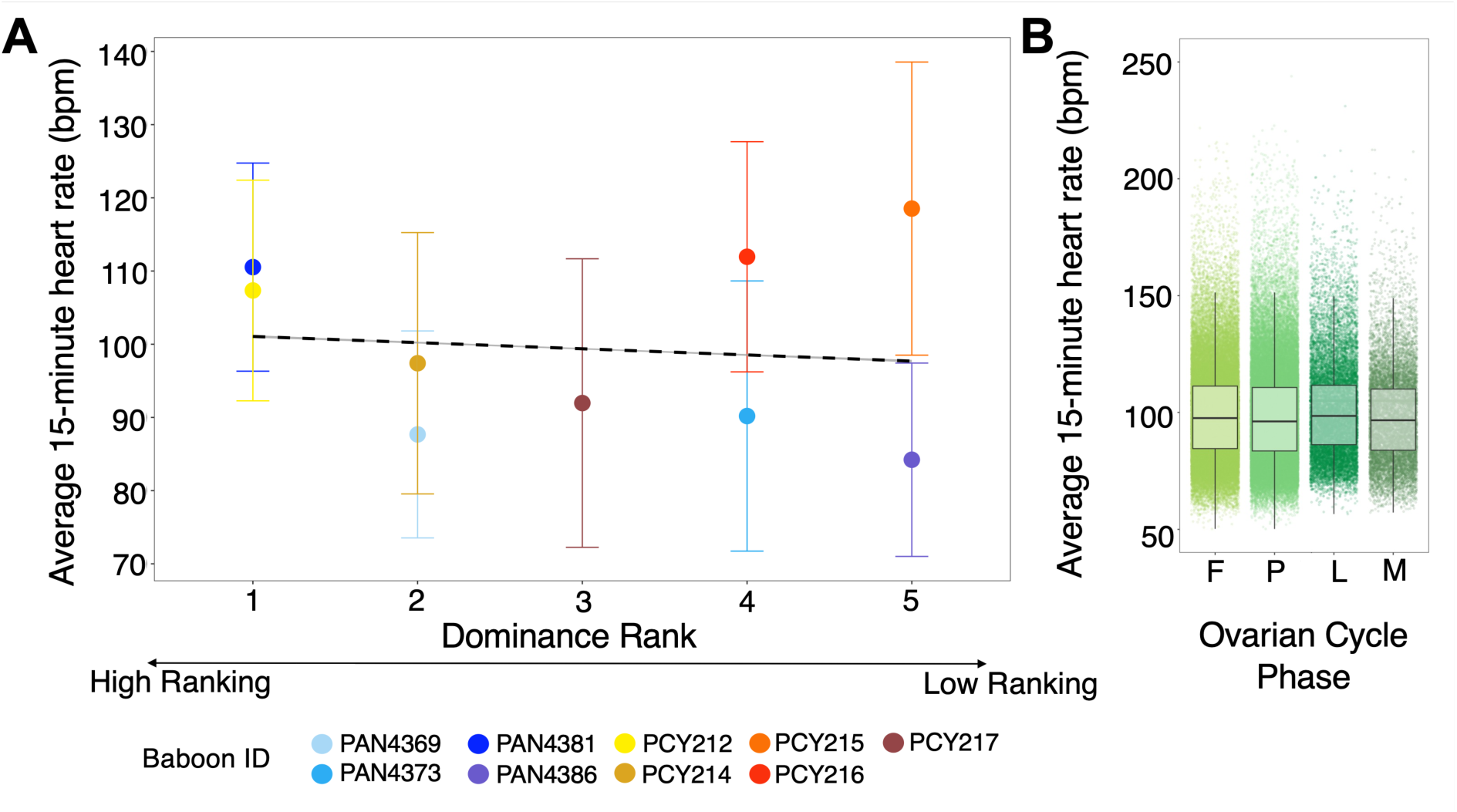
Dominance rank and ovarian cycle phase predict baboon heart rate. Plots show the relationship between 15-minute average heart rate over the 24-hour period (y-axes) and either (**A**) dominance rank or (**B**) ovarian cycle phase. The dashed line in (**A**) shows the linear correlation between baboons’ average 15-minute heart rate and dominance rank. Plot (**B**) shows variation in average 15-minute heart rate (y-axis) as a function of ovarian cycle phase (x-axis). Letters on the x-axis represent different ovarian cycle phases (F = follicular, P = periovulatory, L = luteal, M = menstruation). The bottom and top whiskers represent the minimum and maximum average 15-minute heart rate values respectively. The bottom edge of the box and the top edge represent the 25^th^ and 75^th^ percentile respectively. Blue to purple points represent *P. anubis* individuals and yellow to red points represent *P. cynocephalus* individuals. Shades of green represent different ovarian cycle phases.

### Objective 3: Predictors of physical activity levels in captive female baboons

The best-supported model was the most complex model (ι1AIC >2 compared to all other models; **Table 4**; **Table S3**). Baboon identity explained 2% of the variation in the number of active minutes in a 15-minute period. Controlling for this variation, the number of active minutes was also predicted by body mass, implant location, baboon species, ambient temperature, whether the measures were collected during the day or at night, dominance rank, and ovarian cycle phase (**Table 4**).

**Table 4.** Summary of the best-supported LMM (ACT9 in **Table S3**) explaining variation in activity levels in female baboons at KIPRE. This model was >2 AIC units from all other models (**Table S3**). Reference categories for each fixed effect are denoted in parentheses.

| *Fixed effects* | *Estimate (± SE)* | *T-value* |
| --- | --- | --- |
| *Species (P. cynocephalus)* | -1.62 (0.23) | -7.18 |
| *ICM implant location (pectoral)* | 0.14 (0.25) | 0.57 |
| *Body mass (kg)* | -0.13 (0.01) | -19.09 |
| *Day or night* | -2.63 (0.01) | -197.48 |
| *Ambient temperature ($^{\circ}$C)* | -0.04 (0.00) | -28.48 |
| *Dominance rank* | -0.02 (0.08) | -2.57 |
| *Cycle phase: Periovulatory (Follicular)* | -0.11 (0.01) | -8.36 |
| *Cycle phase: Luteal (Follicular)* | -0.07 (0.02) | -3.81 |
| *Cycle phase: Menstruation (Follicular)* | 0.07 (0.02) | 3.35 |

The largest changes in activity levels were connected to time of day (**Table 4**; individuals were active for nearly 3 minutes more in a given 15-minute period during the day) and baboon species (**Table 4**; *P. cynocephalus* individuals were typically less active by nearly 1.6 minutes in a 15-minute period compared to *P. anubis* individuals). Activity peaked during daylight hours and was more variable during the day than at night (**Figure 4B**). The lowest activity levels were recorded between 23:00 to 06:00, before increasing at 06:00 and often reaching peak levels between 09:00 and 10:00 (**Figure 4B**). Activity levels typically dipped in the middle of the day, peaking again around 16:00 and decreasing after 18:00. Activity levels also varied by species, particularly during daylight hours compared to nighttime hours (**Figure S9A**). *P. cynocephalus* individuals were active for an average of 1.98 minutes (SD ±2.29 minutes) in a 15-minute period during daylight hours compared to *P. anubis* individuals who were active for an average of 3.90 minutes (SD ±3.48 minutes) in a 15-minute period during daylight hours.

While the effect was small, high-ranking animals were less active than low-ranking animals (**Table 4; Figure S9B**). With respect to ovarian cycle phase, females were less active during the periovulatory and luteal phases, and more active during the menstrual phase, compared to the follicular phase (**Table 4; Figure S9C**). The baboons were also more active when temperatures were warm (**Figure S10**).

## DISCUSSION

Implantable cardiac monitors (ICMs) are providing powerful new insights into the physiology and lived experience of animals, with implications for animal health, cognition, welfare, and conservation (14,23,28,31,34,37,38,61). These devices must be tested for safety and accuracy before using them to gain insight into more complex physiological questions; here we find that the Reveal LINQ^TM^ ICM is safe for baboons and provides biologically valid measures of heart rate and physical activity. Our results also contribute new data on heart rate, physical activity, and body temperature for unsedated, unrestrained baboons living in captivity. In addition, we identify several aspects of the baboons’ external environments and internal states that predicted variation in heart rate and physical activity levels, including circadian patterns and associations with dominance rank and ovarian cycle phase. Our study contributes to the growing body of physiologger studies in animals and lays the groundwork to use ICMs in larger scale projects in baboons. Below, we discuss the implications of our results as they pertain to our three objectives.

### ICM safety, data duration, and accuracy

Our procedure for deploying ICMs was similar to that used in prior studies using Medtronic ICMs in animals, including research on American black bears (*Ursus americanus*), maned wolves (*Chrysocyon brachyurus*), scimitar horned oryx (*Oryx dammah*), great apes (*Gorilla gorilla gorilla, Pan troglodytes, and Pongo abelii*), and Gelada baboons (*Theropithecus gelada)* 6/22/26 12:09:00 PM. Similar to these studies, the baboons in our study had no adverse reactions to the implant, however, one implant was rejected by one baboon about a month after deployment. This rate of rejection is lower than in the studies listed above that reported rejection rates. We believe the rejection we observed may have been caused by an incision pocket that was too shallow. As a result, the ICM may have rested close to the incision site, increasing the likelihood that the ICM would migrate out of the incision or be manually removed by the baboon. In other animal studies, rejections typically coincide with observations of infection (37,39), which we did not observe. In humans, when Reveal LINQ^TM^ ICMs are rejected without signs of infection, the rejection is typically attributed to shallow implant pockets, erosion of the skin (loss of the skin’s epithelium, exposing the device and causing it to migrate out of the incision site), or spontaneous device extrusion (62–64). Similar to our study, rejections of this nature in humans also typically occur within a month of implantation. However, we are unable to assign a definite cause to the case of ICM rejection we observed without more detailed observations of this individual.

In addition to assessing the safety of the Reveal LINQ^TM^ in baboons, we were also interested in the device’s longevity in our system. In our study, the Reveal LINQ^TM^ had an average battery life of 191 days. At nominal settings (i.e., the ICM is not programmed with B-ware software and transmits to the MyCareLink^TM^ network once every 24 hours), the Reveal LINQ^TM^ is projected to have a battery life of up to 3 years. By contrast, prior studies using LINQ ICMs reported longer battery life than our study: one year in maned wolves, 16-21 months in scimitar horned oryx, and 1-3 years in other non-human primates (14,31,37,38). We believe that the relatively short lifespan of the Reveal LINQ^TM^ in our study was caused by our high rate of data transmission to the MyCareLink^TM^ network (i.e., hourly instead of a more standard rate of once per day) and how frequently we downloaded B-ware data using the Medtronic 2090 programmer (i.e., monthly instead of >6 months). Establishing repeated successful telemetry connections can increase the device’s energy use and result in faster battery depletion (65,66). The technical difficulty we encountered with the device implanted in PAN4372 that prevented us from downloading her data collected by B-Ware has not been described in other studies where similar biologgers were used in wild species.

In terms of data accuracy, the Reveal LINQ^TM^’s algorithm for measuring R-wave intervals was designed for human clinical use, and differences human and baboon cardiac and musculoskeletal anatomy and physiology could produce variation in electrical noise, resulting in the detection of spurious heart beats (36). Hence it is important to test that the Reveal LINQ^TM^’s heart rate recordings are accurate for baboons. We found that the Reveal LINQ^TM^ provides accurate data on heart rate for baboons: the ICM detected R-waves with clean signal and strong amplitudes that were high relative to the P-, and T-wave amplitudes. For the Reveal LINQs, we considered an R-wave amplitude of 0.15 mV as indicative of reliable ECG sensing, and the mean R-wave amplitude across all individuals in our study was 0.29 mV. We found larger P- and T-wave amplitudes in *P. cynocephalus* as compared to *P. anubis*. This difference may be due to differences in body size and anatomy between species *P. cynocephalus* are taller and thinner than *P. anubis,* who are shorter, stockier, and deep chested (67–69).The comparatively narrower chests of *P. cynocephalus* individuals may have allowed for closer placement of the Reveal LINQ^TM^ to the heart, allowing for better sensing of P- and T-wave amplitudes compared to *P. anubis* individuals. We found that the Reveal LINQ^TM^ sometimes over sensed heart beats when it was placed laterally rather than pectorally in the parasternal area. Specifically, in lateral placements, the ICM was farther from the heart and more subject to musculoskeletal noise produced by arm movements compared to pectoral placements. This musculoskeletal noise was evident when analyzing ECG strips from laterally placed devices. Hence, we recommend pectoral placement for Reveal LINQ^TM^ ICMs in future studies of baboons.

### Physiological parameters for captive female baboons

To our knowledge, our study is one of the first to report heart rate for *P. anubis* and *P. cynocephalus* in unsedated and unrestrained animals. Prior to our study, researchers measured baboon heart rate in baboons that were sedated, restrained, or wearing an ECG tethered vest (41,70–72). Overall, we found lower mean heart rates than these prior studies, with daytime mean 2-minute heart rate ranging from 89.80 – 128.10 bpm, and nighttime mean 2-minute heart rate ranging from 79.0 - 108.80 bpm across subjects. By comparison, Kaminer (70) measured heart rate in 15 sedated chacma baboons (*P. ursinus*) and found a mean heart rate of 122 bpm, ranging from 100-144 bpm. Further, in 170 sedated, captive *Papio* sp. individuals, mean heart rate was 125 bpm (SD ±30 bpm) (71). In unsedated, restrained *P. anubis* and *P. cynocephalus*, mean heart rate was 230 bpm (SD ±46 bpm) (72). The closest values to our study were recorded in unsedated animals tethered and wearing an ECG vest; in these animals mean heart rate was 107.90 bpm (21). While our data encompassed all of these values, because of the relatively non-invasive nature of the ICM, we were able to capture lower, and likely more naturalistic values for mean heart rate in these species.

Our findings on activity patterns in captive baboons mirror those found in other baboons in captivity or the wild (73–76). Similar to wild baboons, whose activity is largely dictated by foraging needs (73,74), the baboons at KIPRE seemed to be influenced by feeding times. Activity levels increased in morning hours after descent from sleeping sites and peaked between 09:00 to 10:00 when the animals were fed, in the mid-afternoon when baboons were provided supplementation of fresh produce, and again before ascent to their sleeping sites in the evening. Similar patterns have been observed in wild baboons (73,74), however wild baboons on average spend a larger part of their day active compared to captive baboons due to constrained resources and increased threats (i.e. predation). However, it is important to note that the Reveal LINQ^TM^’s algorithm to measure physical activity levels cannot be calibrated to a level of sensitivity that is specific to baboons movements. Hence, the Reveal LINQ^TM^ likely did not capture all movements of the baboons in our study. Finally, we found our data on body temperature was consistent for other reports of body temperature for healthy, non-febrile baboons (77,78).

### Predictors of heart rate and activity in captive female baboons

Individual identity was by far the strongest predictor of heart rate in our subjects, but this was not the case for activity levels. Hence, the variation we observed among individual baboons’ cardiac physiology is likely linked to other factors besides their physical activity, including differences in individual reactivity, perception of environmental events, personality, metabolic rates, genetics, and other factors. Individual differences in cardiac physiology have been reported where cardiac data was collected longitudinally in several other unsedated and free-moving individuals of different animal species, including grey seals (*Halichoerus grypus*), Przewalski’s horses (*Equus ferus przewalski*), and yellow-eyed penguins (*Megadyptes antipodes*) (79–82). For example, in yellow-eyed penguins, individuals varied in their responses to human-caused environmental perturbations (82). While we found that individual activity levels were not especially personalized, activity was a reliable predictor of heart rate and showed a strong circadian pattern typical of a diurnal species. Activity patterns were low during nighttime hours due to the physical quiescence of sleep and the dominance of the parasympathetic branch of the nervous system (“rest and digest”). During the day, heart rate was higher and more variable, due to schedules of feeding and animal care and opportunities for social interactions. Heart rate was also predicted by ambient temperature with higher activity and heart rate when ambient temperatures were warmer.

We found that dominance rank had a detectable but small effect on heart rate and physical activity. High ranking animals had higher mean heart rates and were more active than low-ranking animals, controlling for time of day, temperature, body mass, and other variables. The effect of dominance hierarchies on physiology has been widely studied due to the stress associated with different rank positions (1,16,43,44,83). These effects can vary by species, sex, and time scale (83). But, in the “despotic” dominance hierarchies experienced by female *P. anubis* and *P. cynocephalus*, low-ranking individuals are thought to experience lower access to resources and higher rates of aggression than high-ranking individuals (84). Without more detailed behavioral data, it is difficult to know why the high-ranking animals in our study tended to have higher heart rates and activity levels, but these effects could be due to rank-related differences in social behavior or metabolic rates. Because heart rate is a proxy for metabolism (22,85), lower heart rates may be related to lower metabolic rates (86). Lower metabolic rates in low-ranking individuals may be an adaptation to cope with restricted resource access as a function of their position in the hierarchy (87). In contrast, high-ranking individuals may have higher metabolic rates because they are able to take in more energy and may engage in higher rates of energetically costly behaviors, such as mating and aggression than lower ranking individuals (88). In support of this hypothesis, higher metabolic rates in high-ranking individuals have been observed in some species of fish, mammals, and birds (79,88,89). For example, in female red deer (*Cervus elaphus),* low-ranking females have lower resting heart rates than high-ranking females and lost less mass when experiencing periods of food restriction compared to high-ranking individuals (89). In wild baboons, high-ranking females have been observed to have greater food intake and better access to food resources (90,91). However, top-ranked alpha female baboons have lower glucocorticoid levels than other females, suggesting that, in the wild, the effects of rank on metabolic rate may differ from the patterns in captivity (48). Further work is needed to understand rank-related differences in heart rate and activity in baboons.

While the effect was small, we also found that mean heart rate varied as a function of ovarian cycle phase, controlling for other variables. Compared to the follicular phase, individuals had higher average 15-minute heart rates in the periovulatory phase and the luteal phase and lower average 15-minute heart rates during menstruation. Ovarian cycle phase is linked to well-documented changes in hormone levels in humans and other primates, including baboons (92–94). In humans, some studies suggest that changes in these hormone levels may be linked to changes in autonomic nervous system function over course of the ovarian cycle (52,95–97). Previous studies in human females show that regions in the brain and spinal cord that are related to cardiovascular function possess receptors for steroid hormones such as estrogen and progesterone, both of which are associated with ovarian cycling (98–100). Hence, steroid hormones associated with female reproduction could directly influence cardiovascular functioning, but this connection is tenuous. It is also possible that cycling female baboons experience changes in activity levels and social behavior as they transition through the different ovarian cycle phases. For example, female baboons may become more susceptible to male and female aggression as they enter their periovulatory period as a result of reproductive competition (101–103). Changes in activity levels over the ovarian cycle could be connected to changes in heart rate but we did not observe that activity was strongly predicted by ovarian cycle phase in our study. Given that baboons serve as a strong model for human female reproduction, investigating such a question may aid in understanding physiological variation that coincides with female reproductive changes.

## CONCLUSIONS

We suggest three main directions for future work. First, it will be valuable to develop protocols to download data from LINQ ICMs remotely, without the need to re-capture and sedate the animal. This ability may be enabled by Bluetooth signals produced by newer models of the LINQ and the potential to transmit data to cloud-based repositories over cellular or satellite networks (18). Second, such enhanced download capability of ICMs could provide a new perspective on the nexus between social behavior, physiology, health, and survival in wild populations of baboons. Natural populations of baboons have made valuable contributions to elucidating these connections (45,102,104–106), revealing links between social rank and health (43,47,48,83), or between early-life adversity and longevity (107,108). However, testing the physiological mechanisms that underlie these effects has been challenging. To date, many of these insights have come from data on non-invasively sampled glucocorticoid metabolites (45,46,106,109) that provide a relatively coarse perspective on stress responses and energy mobilization. The ICMs we test here could provide fine-grained data to understand how natural social and environmental events “get under the skin” to affect individual health and survival.

Third, given the strong circadian patterns in heart rate and activity that the Reveal LINQ^TM^ ICMs detected, deploying ICMs in wild baboons may also provide insight into sleep physiology and behavior and how sleep phenotypes vary between individuals. Behavioral observations of sleep-related behaviors in animals have yielded significant descriptive findings on sleep ecology within populations and across species (110). However, without detailed physiological data, it is difficult to ascertain how factors related to sleep quality and duration vary due to biological, ecological, and social factors.

In summary, our work represents a useful next step in expanding research on cardiac physiology in freely moving animals. Such advances are essential to learn how individuals cope with internal and external environmental changes in natural contexts. We hope that our work will aid in the development of new methods to measure individual behavior, stress responses, metabolism, and circadian rhythm of animals in more naturalistic contexts. Furthermore, given that cardiac parameters have been connected to the development of chronic diseases and longevity outcomes (111–113), our work represents a valuable springboard for understanding sources of variation in the development of patterns of aging, disease, and mortality in wild species.

## LIST OF ABBREVIATIONS

ICM: insertable cardiac monitor
HR: heart rate
Bpm: beats per minute
ECG: electrocardiogram
KIPRE: Kenya Institute of Primate Research

## DECLARATIONS

### Ethics Approval

All research was approved by the Institutional Animal Care and Use Committee (IACUC) at the University of Notre Dame and the Institutional Scientific and Ethics Review Committee at KIPRE (ISERC/09/22); all research also adhered to the laws and guidelines of the Kenyan government (NACOSTI permit number: NACOSTI/P/23/22601).

### Consent for Publication

Not applicable.

### Availability of data and materials

The datasets and code supporting the conclusions of this article are available on GitHub (https://github.com/catherine-andreadis/determinants_HR_activity_captiveBaboons.) For peer review, data and code are available at: https://notredame.box.com/s/gsnxji4la1lybjquw0ja0ud88j1s23ah.

### Authors’ contributions

CRA, IGK, HP, TGL, MYA, and EAA conceptualized the study. CRA, IGK, JN, DK, PK, KMK, NRL, JM, NW, TGL, MYA, and EAA contributed to the investigation and data collection. CRA performed the statistical analyses and data visualizations. CRA and EAA wrote and prepared the original manuscript draft.

CRA, IGK, JN, DK, PK, KMK, NRL, JM, NW, HP, TGL, MYA, and EAA reviewed and edited the manuscript. HP, TGL, MYA, and EAA acquired the funding.

## Acknowledgements

We thank the University of Notre Dame and the Welfare and Ethics Directorate of the Kenya Institute of Primate Research (KIPRE) for logistical support. We also thank members of KIPRE colony management and primate staff. In Kenya, our research was approved by the National Commission for Science, Technology, and Innovation, and the National Environment Management Authority. All ICMs and monitors were donated by Medtronic.

