## Supplemental Materials for "Using insertable cardiac monitors to test determinants of heart rate and activity in captive baboons"

### Supplementary Information

### Supplementary Tables

**Table S1.** Welch's Two-sample T-test results

| *Welch's two-sample T-test* | *Mean amplitude in mV (± SE)* | *t* | *p* |
|---|---|---|---|
| *R-wave amplitude (PAN) v. R-wave amplitude (PCY)* | R-wave PAN = 0.271 (0.113)<br>R-wave PCY = 0.335 (0.199) | -1.99 | 0.051 |
| *P-wave amplitude (PAN) v. P-wave amplitude (PCY)* | P-wave PAN = 0.026 (0.023)<br>P-wave PCY = 0.046 (0.045) | -2.77 | 0.007 |
| *T-wave amplitude (PAN) v. T-wave amplitude (PCY)* | T-wave PAN = 0.042 (0.042)<br>T-wave PCY = 0.797 (0.121) | -2.08 | 0.041 |
| *R-wave amplitude (pectoral) v. R-wave amplitude (lateral)* | R-wave pectoral = 0.302 (0.095)<br>R-wave lateral = 0.304 (0.234) | 0.043 | 0.966 |
| *P-wave amplitude (pectoral) v. P-wave amplitude (lateral)* | P-wave pectoral = 0.037 (0.031)<br>P-wave lateral = 0.035 (0.045) | -0.156 | 0.879 |
| *T-wave amplitude (pectoral) v. T-wave amplitude (lateral)* | T-wave pectoral = 0.05 (0.054)<br>T-wave lateral = 0.077 (0.129) | 1.21 | 0.232 |

Welch's Two-sample T-test results for comparisons of R, T, and P ECG wave amplitudes between species (*P. anubis*, PAN or *P. cynocephalus*, PCY, upper table) or ICM implantation sites (pectoral or lateral, lower table).

**Table S2.** AIC results for determinants of heart rate

| *Model* | *Variables included in the model* | *AIC* | *∆AIC* |
|---|---|---|---|
| *HR9* | average 15-minute heart rate ~ active minutes + ambient temperature + day or night + body mass + implant location + baboon species + dominance rank + ovarian cycle phase + (1\|baboon identity) | 1255834.0 | 0.00 |
| *HR8* | average 15-minute heart rate ~ active minutes + ambient temperature + day or night + body mass + implant location + baboon species + dominance rank + (1\|baboon identity) | 1256026.0 | 192.40 |
| *HR7* | average 15-minute heart rate ~ active minutes + ambient temperature + day or night + body mass + implant location + baboon species + (1\|baboon identity) | 1256029.0 | 195.18 |
| *HR6* | average 15-minute heart rate ~ active minutes + ambient temperature + day or night + body mass + implant location + (1\|baboon identity) | 1256034.0 | 200.42 |
| *HR4* | average 15-minute heart rate ~ active minutes + ambient temperature + day or night + (1\|baboon identity) | 1256484.0 | 649.75 |
| *HR5* | average 15-minute heart rate ~ active minutes + day or night + (1\|baboon identity) | 1257764.0 | 1929.72 |
| *HR3* | average 15-minute heart rate ~ active minutes + ambient temperature + (1\|baboon identity) | 1266338.0 | 10503.70 |
| *HR2* | average 15-minute heart rate ~ active minutes + (1\|baboon identity) | 1274840.0 | 19005.83 |
| *HR1* | average 15-minute hear rate ~ (1\|baboon identity) | 1285570.0 | 29735.75 |

The nine best-supported models explaining variation in 15-minute average HR based on AIC and delta AIC (∆AIC).

**Table S3.** AIC results for determinants of activity levels

| *Model* | *Variables included in the model* | *AIC* | *∆AIC* |
|---|---|---|---|
| *ACT9* | active minutes ~ ambient temperature + day or night + body mass + implant location + baboon species + dominance rank + ovarian cycle phase + (1\|baboon identity) | 673062.4 | 0.00 |
| *ACT7* | active minutes ~ ambient temperature + day or night + body mass + implant location + baboon species + (1\|baboon identity) | 673147.4 | 85.05 |
| *ACT8* | active minutes ~ ambient temperature + day or night + body mass + implant location + baboon species + dominance rank + (1\|baboon identity) | 673147.8 | 85.41 |
| *ACT5* | active minutes ~ ambient temperature + day or night + body mass + (1\|baboon identity) | 673155.7 | 93.35 |
| *ACT6* | active minutes ~ ambient temperature + day or night + body mass + implant location + (1\|baboon identity) | 673156.6 | 94.19 |
| *ACT3* | active minutes ~ ambient temperature + day or night + (1\|baboon identity) | 673491.2 | 428.80 |
| *ACT4* | active minutes ~ day or night + (1\|baboon identity) | 674090.2 | 1027.83 |
| *ACT2* | active minutes ~ ambient temperature + (1\|baboon identity) | 707828.8 | 34766.47 |
| *ACT1* | active minutes ~ (1\|baboon identity) | 712784.9 | 40722.55 |

The nine best-supported models explaining variation in 15-minute activity level based on AIC and dela AIC (∆AIC).

### Supplementary Figures

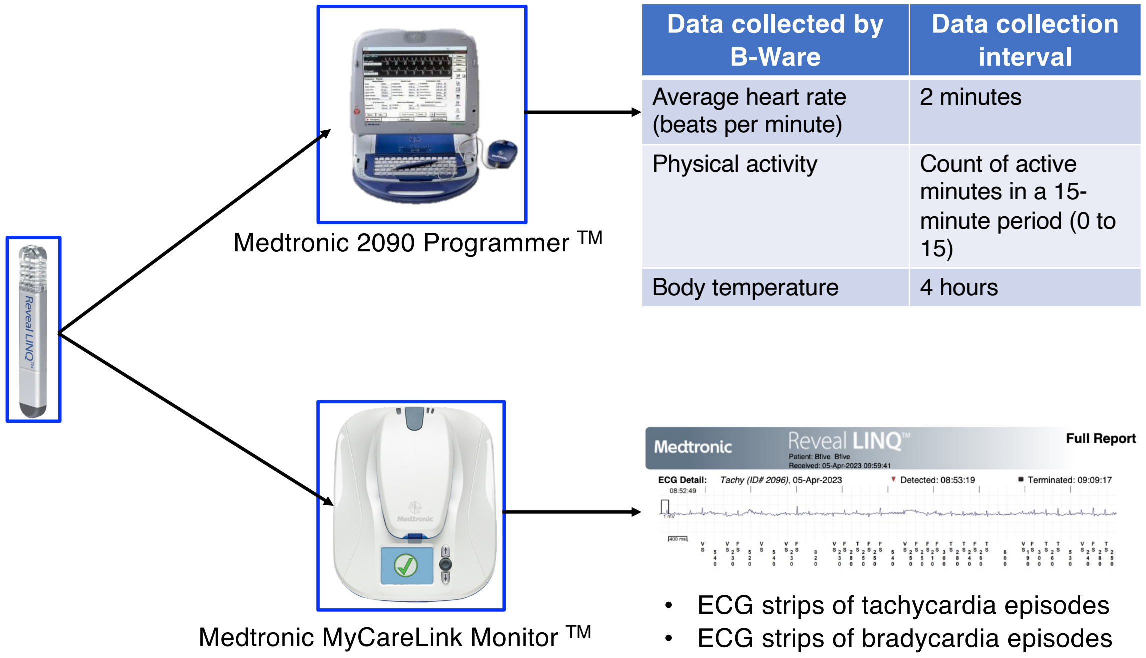

| Data collected by B-Ware | Data collection interval |
|---|---|
| Average heart rate (beats per minute) | 2 minutes |
| Physical activity | Count of active minutes in a 15-minute period (0 to 15) |
| Body temperature | 4 hours |

**Figure S1. Flowchart of equipment used during monthly data downloads from Reveal LINQ™ ICMs.** This flowchart shows the equipment used to download different types of data from the Reveal LINQTM ICMs. The Medtronic 2090 Programmer™ (upper flowchart path) was used to download three data sets collected by B-Ware software shown in the inset table: 2-minute average heart rate, physical activity as a count of active minutes in a 15-minute period, and body temperature every 4 hours. The Medtronic MyCareLink Monitor™ was used to download ECG strips associated with tachycardia and bradycardia episodes (lower flowchart path). These data could then be accessed on the Medtronic CareLink Network website.

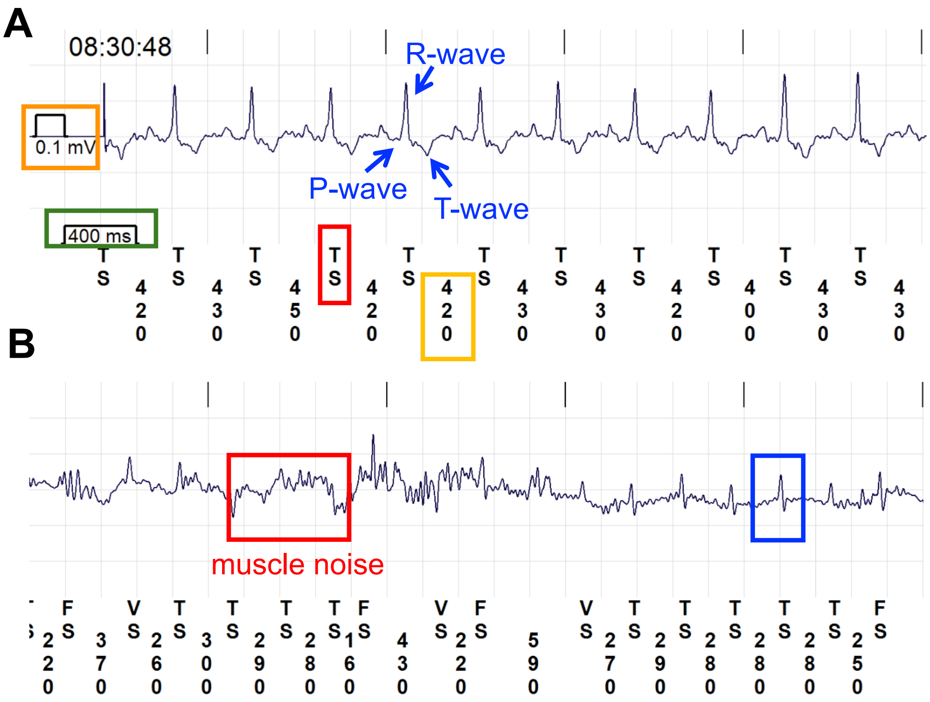

**Figure S2. Two representative ECG strips recorded during tachycardia events for (A) PAN4386 and (B) PAN4373.** Panel **(A)** shows the P-, R-, and T- wave of a discernible ECG waveform marked with blue arrows. The scale for time in milliseconds (ms) is on the x-axis and highlighted in green. The scale for wave amplitude (mV) is shown on the y-axis and is highlighted in orange. The time for a single R-R interval in ms is highlighted in yellow (420 ms). An R-wave that is defined as a true heartbeat recorded by Medtronic is denoted by TS, VS, and FS notation. An example of this is highlighted in red. The letters "TS" below the ECG trace refer to a tachycardia sensing event. This means that the distance between two R-waves met or exceeded the threshold value for a tachycardia beat (we programmed the Reveal LINQ$^{TM}$ to label any beat 130 bpm or higher as a tachycardia event). The letters "VS" below the ECG trace refers to a ventricular event, or an instance of ventricular depolarization occurring during a heartbeat. The notation "FS" refers also refers to a tachycardia event, but unlike "VS" events, the Reveal LINQ$^{TM}$ categorizes these as potential fibrillation events. Fibrillation events differ from tachycardia events in that heart beats during fibrillation events are fast and irregularly timed, while heart beats during tachycardia events are fast and regularly spaced. Panel (**B**) shows a tachycardia episode for PAN4373. Muscle noise, which was collected and identified as heart rate data by the Reveal LINQ$^{TM}$, as evidenced by the assignment of TS notation to these peaks in the red box, is highlighted in red. A correctly sensed heartbeat discerned by a clear ECG waveform is highlighted in blue. Definitions for TS, VS, and FS notation as seen in ECG strip in panel (**B**) are identical to those described for panel (**A**).

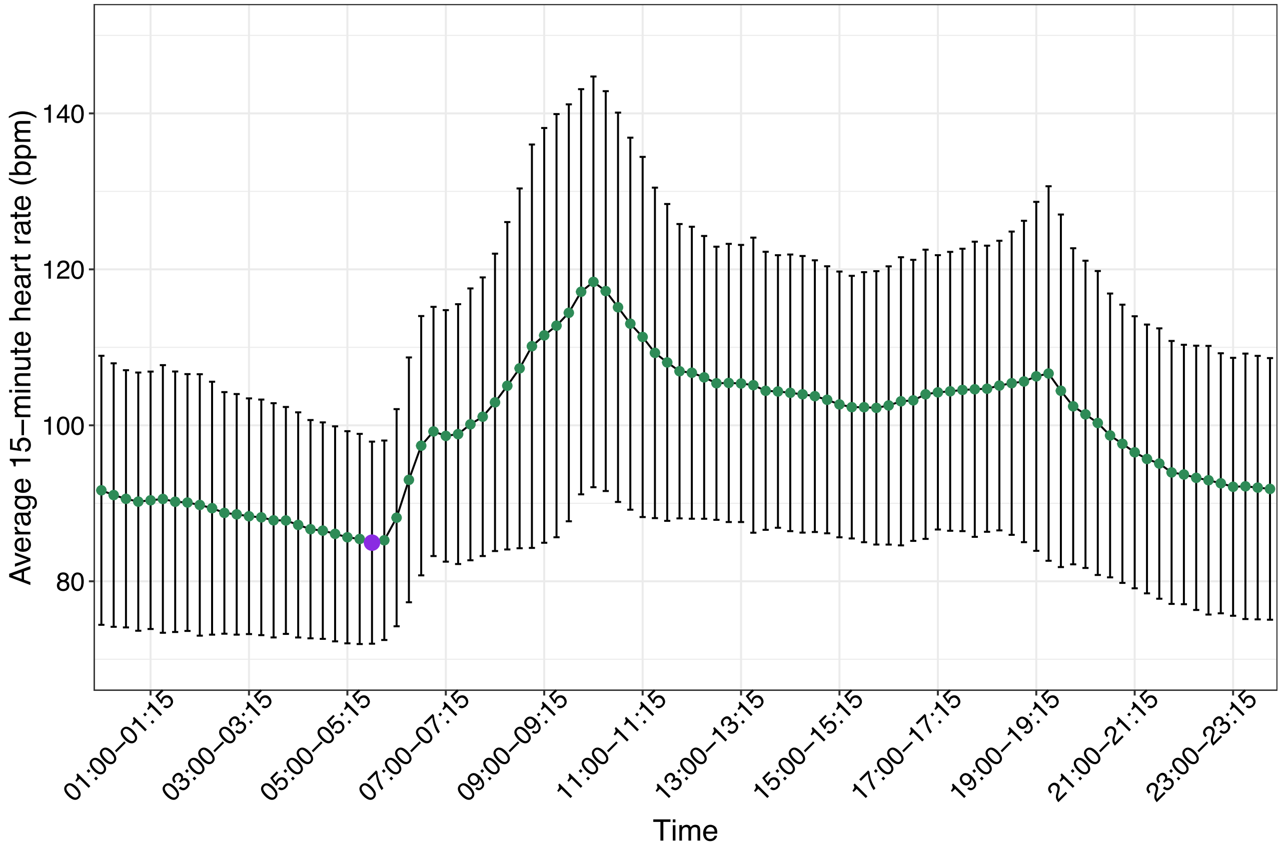

**Figure S3. Baboon heart rate was lowest between 05:30 and 05:45, which was selected as the period for resting heart rate.** Plot depicts the mean 15-minute average heart rate in bpm (y-axis) plotted over 15-minute windows (x-axis). Each point represents the average-15-minute heart rate for a 15-minute window averaged across all 9 baboons. The larger, purple point represents the average 15-minute heart rate for 05:30-05:45. Error bars represent standard error of the mean.

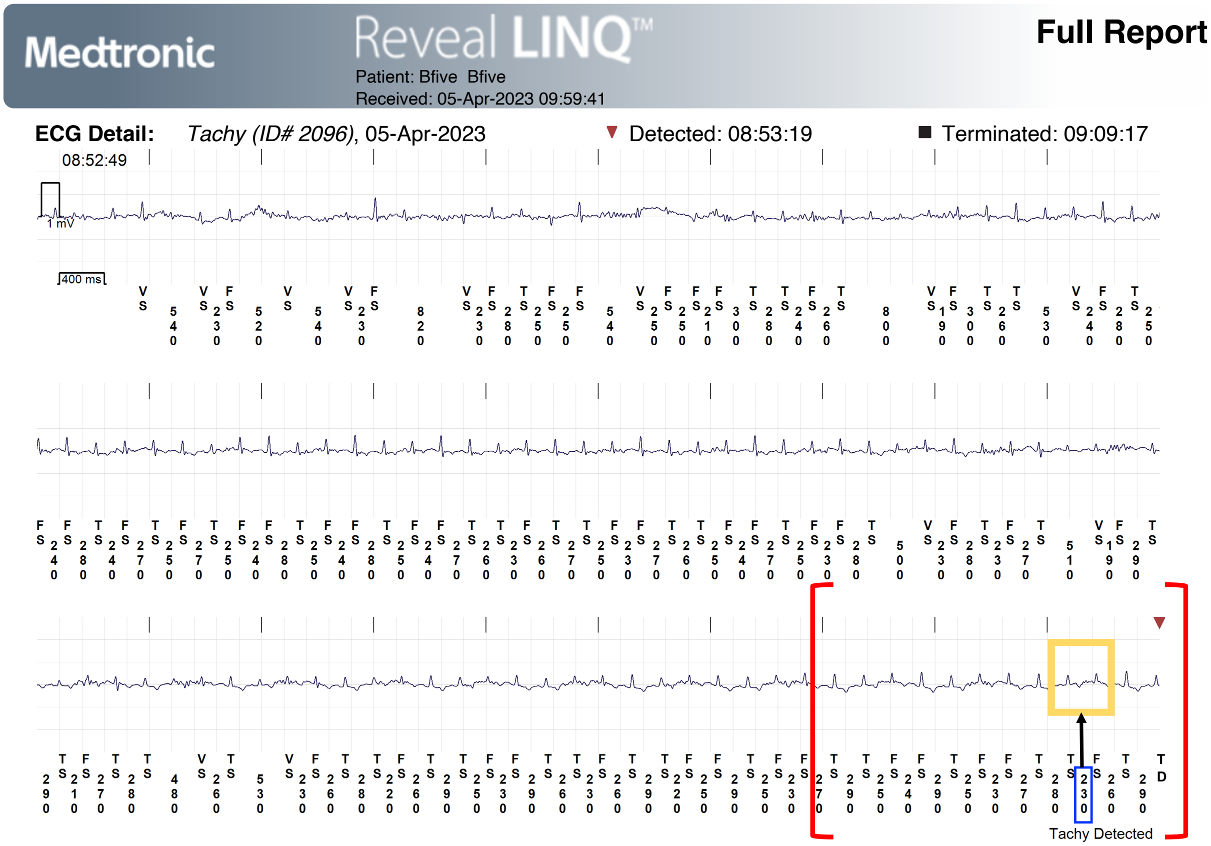

**Figure S4. ECG strip depicting the maximum validated heart rate during a tachycardia episode.** This strip was recorded by the Reveal LINQ$^{TM}$ in baboon PCY214 on 5 April 2023 at 08:53. The red brackets denote the beginning and end of the tachycardia episode (defined as 12 consecutive beats at 130 bpm or higher). The value boxed in blue ("230") refers to a representative R-R interval in milliseconds of the highest validated heart rate of 260 bpm. The mean heart rate across all 12 beats of the bracketed tachycardia episode was 228.6 bpm. The ECG waves boxed in yellow refer to the distance between the R-wave peaks that generated the 230 millisecond (ms) value. TS, VS, and FS notations are defined in the legend of Figure S1.

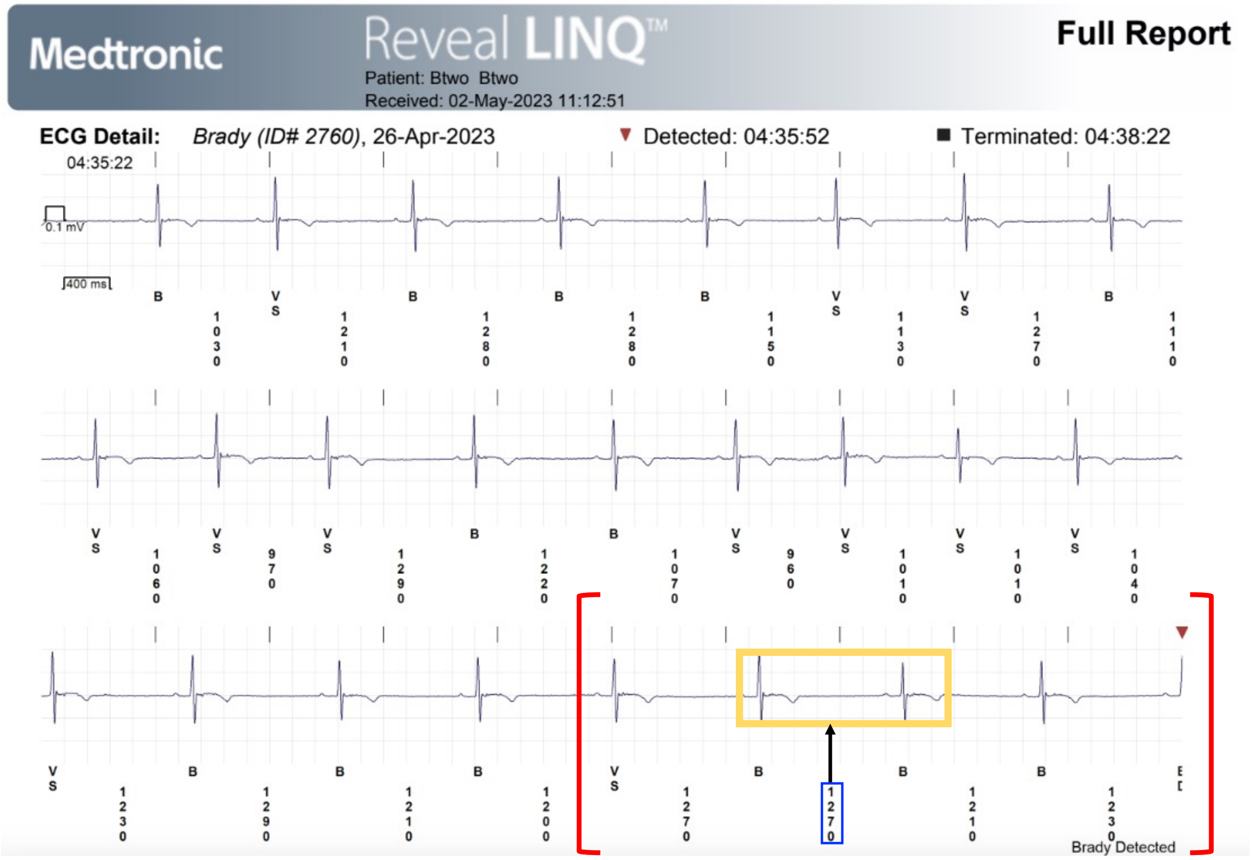

**Figure S5. ECG strip depicting the lowest validated heartbeat during a bradycardia episode.** This strip was recorded by the Reveal LINQ™ in baboon PAN4372 on 26 April 2023 at 04:35. The red brackets denote the beginning and end of the bradycardia episode (defined as 4 consecutive beats at 50 bpm or lower). The mean heart rate across all 4 beats of the bracketed bradycardia episode was 48.1 bpm (mean = 1245 milliseconds (ms) between consecutive R-waves). The value boxed in blue ("1270") refers to a representative R-R interval in milliseconds of the lowest validated heart rate of 47.4 bpm. The ECG waves boxed in yellow refer to the distance between the R-wave peaks that generated the 1270 ms value. Note that a slower HR was observed prior to the Reveal LINQ's detection of the 4-beat bradycardia episode (46.5 bpm or 1290 ms between two consecutive R-waves). VS notations are defined in the legend of Figure S1 and "B" notations refer to a bradycardia sensing event. This means that the distance between two R-waves met or exceeded the threshold value for a bradycardia beat ($\leq$ 50 bpm).

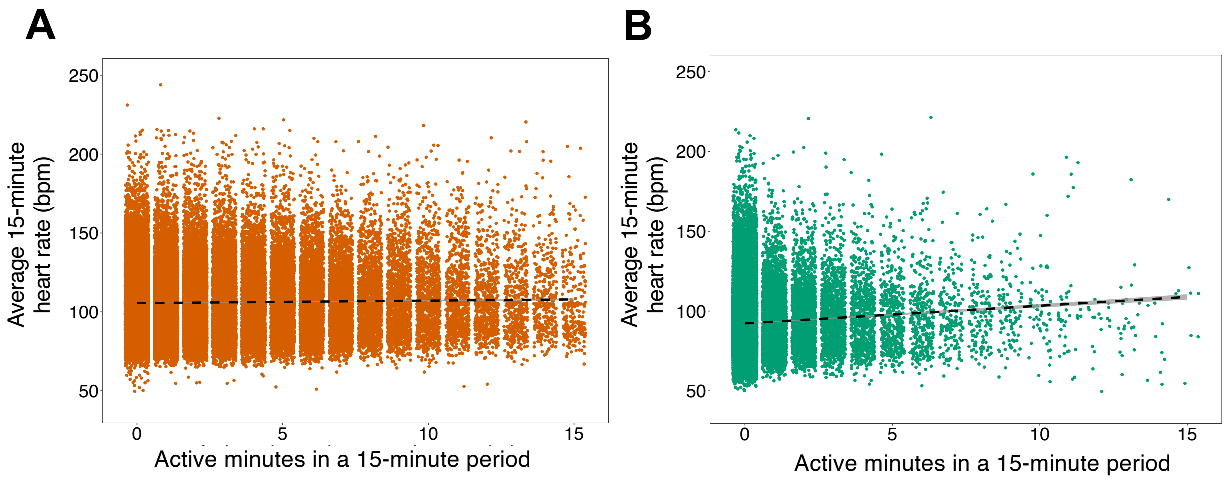

**Figure S6. Baboon heart rate was positively correlated with physical activity.** Plots show the 15-minute average heart rate in bpm (y-axis) as a function of the number of active minutes in a 15-minute period (x-axis) during (A) the daytime (orange) or (B) at nighttime (green; see text for definitions of daytime and nighttime). Dashed lines show linear correlations; grey shaded area around the dashed line shows the 95% confidence interval for the fitted data.

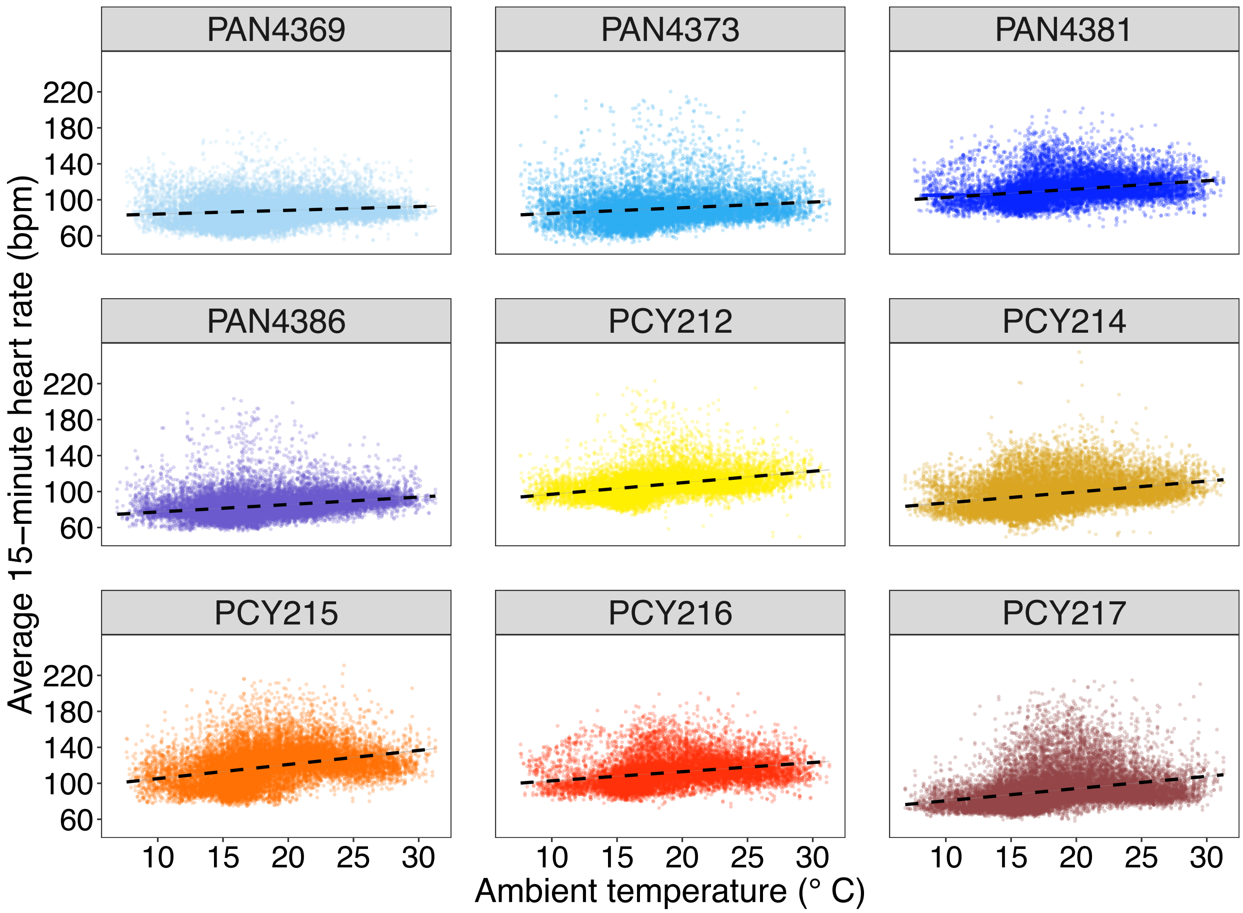

**Figure S7. Baboon average 15-minute heart rate was positively correlated with ambient temperature.** Plots show the relationship between 15-minute average heart rate in bpm (y-axis) and ambient temperature in °C (x-axis) for each individual baboon over the course of data collection (female identity is at the top of each plot). Color gradients represent baboon species; blue to purple points represent *P. anubis* and yellow to red points represent *P. cynocephalus*.

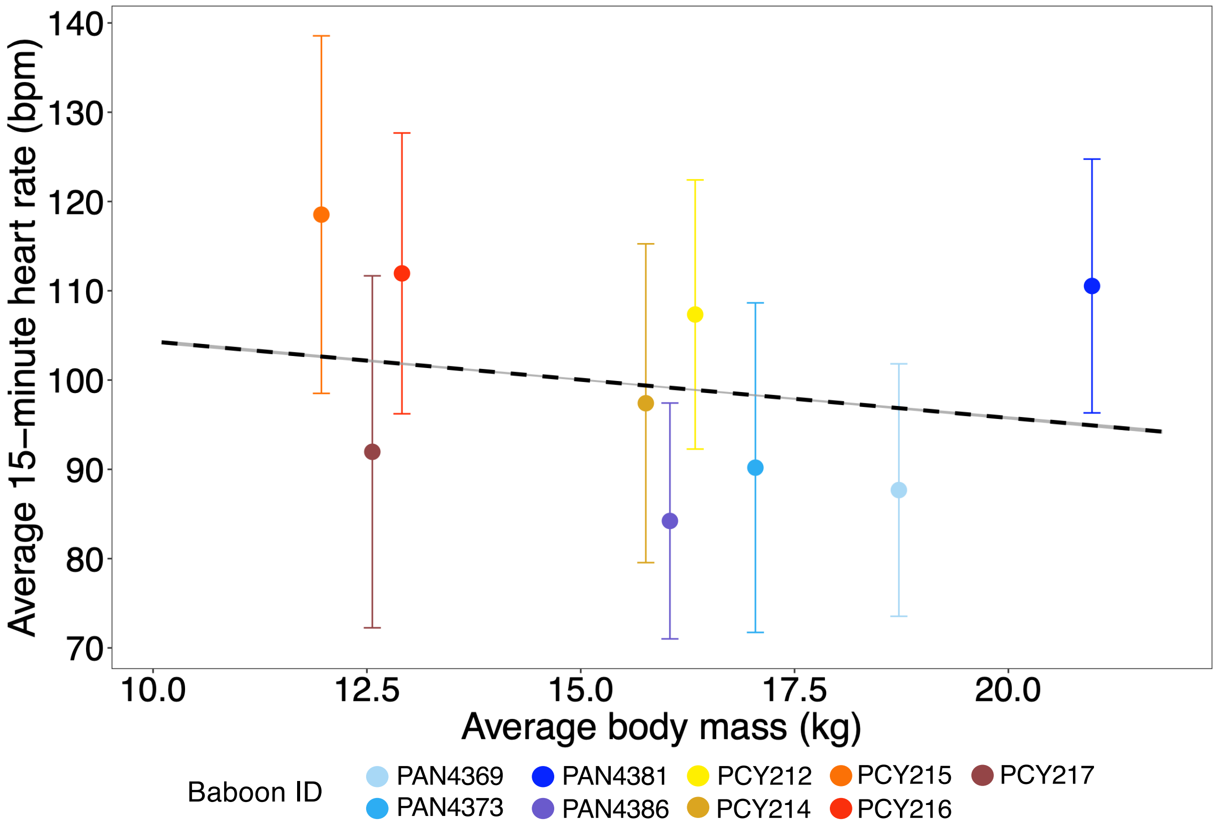

**Figure S8. Individual body mass predicts heart rate.** Plot shows the relationship between 15-minute average heart rate (y-axis) and average body mass (kg) (x-axis). The dashed line shows the linear correlation between baboons' average 15-minute heart rate calculated for the course of the study and average body mass. Color gradients represent baboon species; blue to purple points represent *P. anubis* and yellow to red points represent *P. cynocephalus*.

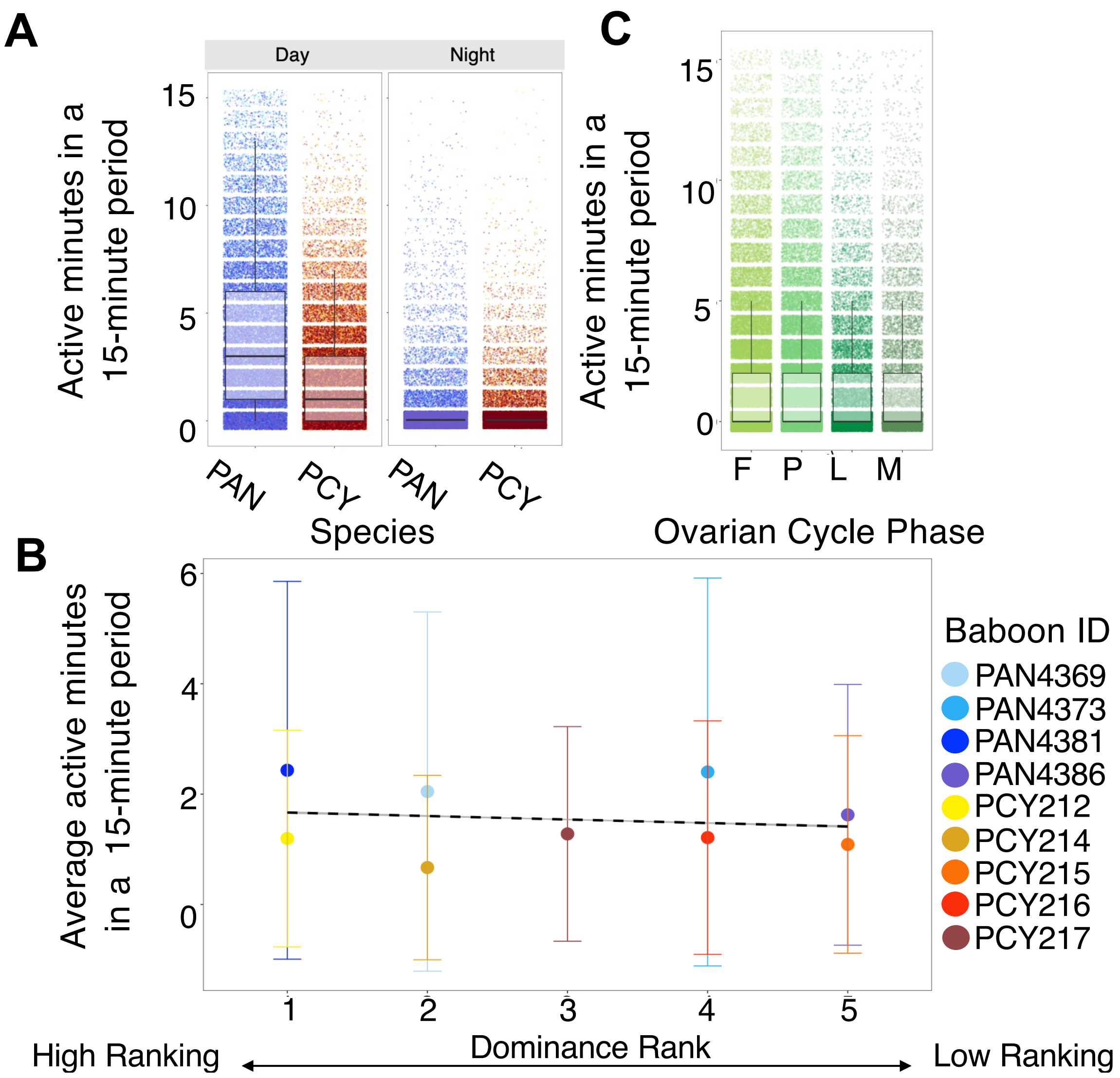

**Figure S9. Species identity, dominance rank, and ovarian cycle phase predict activity levels.** Plots show the relationship between the number of active minutes in a 15-minute period (y-axes) as a function of (**A**) baboon species, (**B**) dominance rank, or (**C**) ovarian cycle phase. In the boxplots in (**A**) and (**C**), the middle line represents the median value for active minutes in a 15-minute period, the bottom and top whiskers represent the minimum and maximum, and the bottom edge of the box and the top edge represent the 25$^{th}$ and 75$^{th}$ quartiles. The dashed line in (**B**) shows the linear correlation between baboons' average activity level in a 15-minute period and dominance rank. Plot (**C**) shows variation in active minutes in a 15-minute period (y-axis) as a function of ovarian cycle phase (x-axis). Letters on the x-axis represent different ovarian cycle phases (F = follicular, P = periovulatory, L = luteal, M = menstruation). In plot (**A**), blue points represent data from *P. anubis* individuals and yellow points represent data from *P. cynocephalus* individuals. In plot (**B**), color gradients represent baboon species; blue to purple points represent *P. anubis* and yellow to red points represent *P. cynocephalus*. In plot (**C**), shades of green represent different ovarian cycle phases.

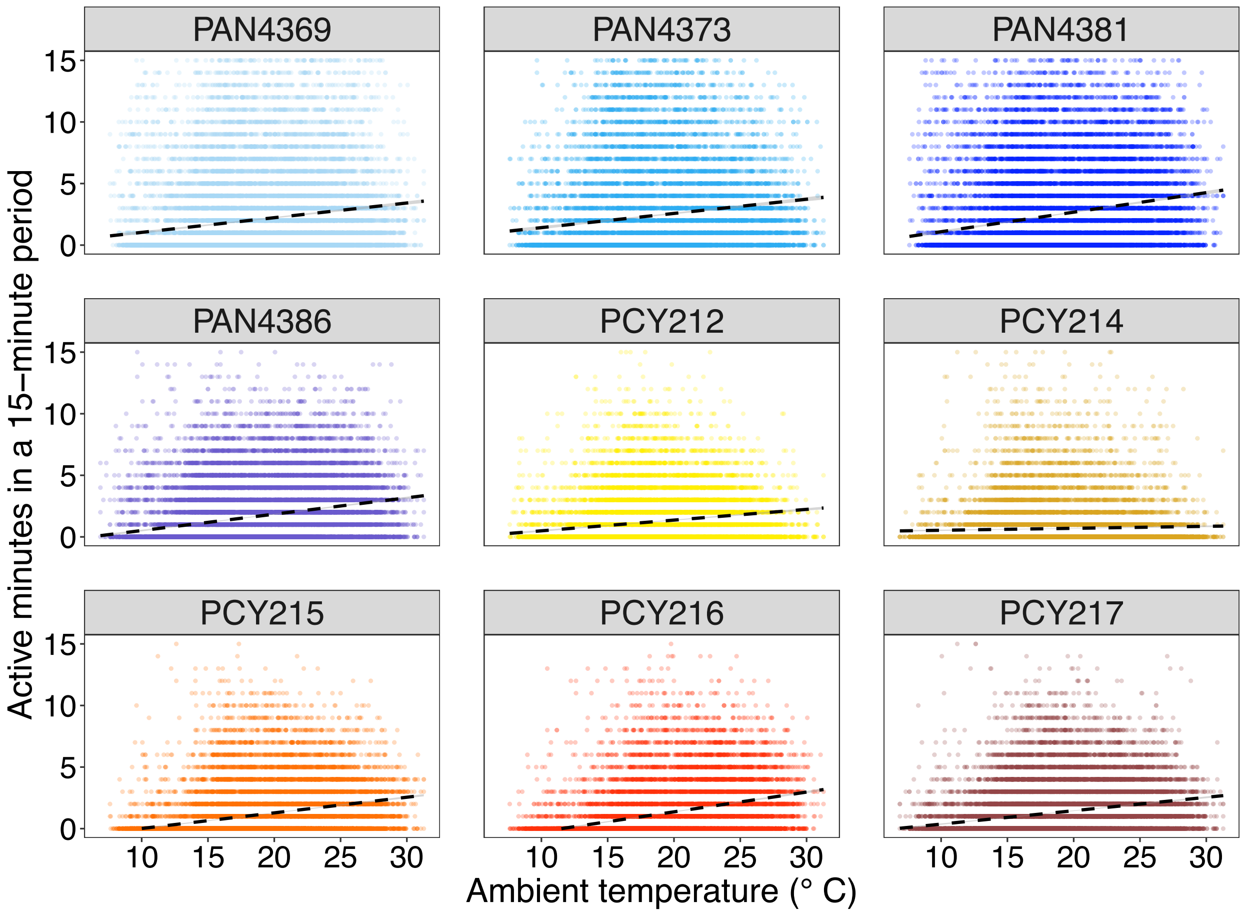

**Figure S10. Active minutes in a 15-minute period was positively correlated with ambient temperature.** Plots show the relationship between active minutes in a 15-minute period (y-axis) and ambient temperature in °C (x-axis) for each individual baboon over the course of data collection (female identity is at the top of each plot). Color gradients represent baboon species; blue to purple points represent *P. anubis* and yellow to red points represent *P. cynocephalus*.
